# Predictive consequences of alternative representations of metabolic measurements: A controlled synthetic study of categories, longitudinal information and transition preservation

**DOI:** 10.64898/2026.10.27.762152

**Authors:** Mwangi Njenga, Waweru Peter

## Abstract

Metabolic categories summarise observed abnormalities, but shared labels may conceal distinctions relevant to future evolution. We examined the predictive consequences of alternative representations under controlled synthetic dynamics.

For each of six generating scenarios, twenty independent confirmation cohorts contained 5,115 simulated individuals each, including 1,023 held out. Observations through year 20 supported year-25 prediction under continued loading or reassignment to a specified low-load regime. Conventional categories, current and longitudinal measurements, selected descriptors and an outcome-guided embedding were evaluated with Ridge regression and histogram-based boosting. Paired cohort comparisons retained their original multiplicity families. A descriptive ablation examined phenotype-bearing and recorded etiological inputs. Separate post hoc work assessed monthly category-transition preservation.

Under scheduled noisy conventional-panel observation and Ridge regression, the selected current and longitudinal representation (Φ) had lower loss than binary syndrome status and criterion count in all twelve scenario–task comparisons against each comparator. In current-state-sufficient continuation, the Φ-minus-binary-status difference was −0.0156 (pointwise 95% confidence interval −0.0187 to −0.0125). No advantage over current-plus-baseline measurements was established under the joint primary correction, and Φ-derived clusters performed better in nonlinear low-load prediction. Stronger categories and different learners altered some comparisons. Adding recorded etiological indicators produced small, context-dependent changes, including adverse increments. Two constructed states sharing syndrome status and count had next-month positive-category probabilities of 10.16% and 69.54%. An independent analytic argument established failure of strong lumpability for that monthly kernel. These were synthetic model probabilities, not patient risks.

Categorisation can conceal future-relevant distinctions without being uniformly inferior for fitted prediction. Performance depends on information access, transformation, task and learner. The findings support explicit evaluation of metabolic representations, but do not establish a predictively sufficient representation of human physiology, clinical replacement of metabolic syndrome or therapeutic benefit.

## Introduction

Metabolic measurements describe observations of physiology that continues to change. A diagnostic category can make those observations useful for communication or decision making without retaining every distinction needed to anticipate their future. Metabolic syndrome illustrates this issue. Its harmonised definition combines waist circumference, blood pressure, glucose, triglycerides and HDL cholesterol, with at least three positive criteria identifying the syndrome (Alberti et al., 2009). Reaven’s physiological account connected several co-occurring abnormalities through insulin resistance, whereas Kahn and colleagues questioned the additional explanatory and prognostic meaning supplied by the syndrome label (Reaven, 1988; Kahn et al., 2005). The association of metabolic syndrome with cardiovascular outcomes establishes clinical relevance, but does not establish that its categories preserve all useful information in their constituent measurements (Mottillo et al., 2010).

Human studies provide direct precedents for examining distinctions within and between categories. Franco and colleagues related component combinations and paths into metabolic syndrome to subsequent cardiovascular disease and mortality (Franco et al., 2009). Baltimore and Whitehall II studies described longitudinal changes in syndrome components and glucose regulation before disease recognition (Scuteri et al., 2009; Tabák et al., 2009). Changes in syndrome status also carried prognostic information in a large national cohort (Park et al., 2019). Continuous severity scores, joint metabolic trajectory models and data-driven clinical groups represent established alternatives to a single binary label (Gurka et al., 2014; Breeze et al., 2016; Ahlqvist et al., 2018).

These approaches answer different questions. Predictive usefulness must be compared with credible alternatives for the specified outcome, as illustrated by comparisons of diabetes subgroups with models using continuous clinical features (Dennis et al., 2019).

Categorisation is a many-to-one transformation: distinct measurement vectors can receive the same label. This loss of distinctions does not determine their importance for a future outcome. A category can discard irrelevant variation or simplify estimation for a finite learner, while a richer representation can introduce poorly estimated or redundant features. Established criticisms of dichotomisation therefore motivate a comparison rather than a presumption that every continuous description must predict better (Altman and Royston, 2006; Royston et al., 2006). Historical access complicates the comparison further. A longitudinal representation may outperform a current category because it receives earlier measurements, because it transforms common information differently, or because the fitted learner uses its inputs more effectively.

A controlled synthetic study can make these distinctions inspectable by separating the state that evolves, the observations generated from it, the information supplied to each representation and the learner making the prediction. Dynamical physiological models and state-and-observation methods provide established foundations for this separation (Topp et al., 2000; Särkkä and Svensson, 2023). Here, METLAS supplies an organising view of metabolic evolution; the evaluated generators implement selected concepts without constituting a numerical validation of the larger framework. Our question is which future-relevant distinctions alternative representations preserve under stated dynamics and observation conditions. We examine conventional and strengthened categories, current and longitudinal measurements, and recorded exposure information. A separate post hoc analysis asks whether categories preserve their own next-month transition probabilities under one identified law. Predictive adequacy, measurement requirements and representation movement are addressed in the companion paper; the present article does not infer those properties from predictive advantage.

## Methods

### Study design and scientific objects

The study used stipulated stochastic metabolic systems rather than human observations. Six scenarios represented current-state-sufficient dynamics (G0), accumulated burden (G1), reserve depletion and recovery (G2), adaptive operating reference (G3), persistent effects of historical exposure (G4), and nonlinear regime-dependent dynamics (G5). These labels describe alternative mathematical mechanisms, not identified human disease subtypes. Four interacting domains connected the latent state to adiposity, glycaemia, lipoprotein metabolism and haemodynamics. Depending on the scenario, present burden was supplemented by cumulative exposure, reserve, adaptation or retained alteration; the nonlinear scenario also included a persistent hidden regime. Table 1 summarises the roles relevant to interpreting the comparisons. The simulation design distinguishes generating mechanisms, estimands and performance measures in the sense described by Morris and colleagues (Morris et al., 2019).

**Table 1.** Synthetic scenarios and the distinctions they make examinable.

| Scenario | Distinction implemented | Interpretive role |
| --- | --- | --- |
| Current-state-sufficient | Present burden and context determine the next-state law | Control for historical benefit without an active historical carrier |
| Accumulated burden | A cumulative-exposure carrier affects future burden | History can influence the transition through a present carrier |
| Reserve depletion and recovery | Reserve changes target and restoration rate | Similar burdens need not have similar responses |
| Adaptive operating reference | A slowly changing reference affects subsequent behaviour | Adaptation persists after input reassignment |
| Persistent historical effects | A slowly reversible alteration affects burden | Earlier exposure can have retained consequences |
| Nonlinear regime-dependent | Hidden persistent regimes and nonlinear interactions | Allows category-favouring prediction without requiring it |

Scenario names describe stipulated mathematical constructions. They are not identified human subtypes. Each scenario contributed twenty independent confirmation cohorts.

The complete generating context included the fixed subject variables and external inputs required by the transition law, whether or not learners observed them. Physiology advanced monthly over 35 years. Specified mappings generated measurements in clinical units; a separate observation process controlled error, visit schedules, missingness, dropout and reporting delays. The conventional panel comprised six measurements producing five diagnostic criteria. Two larger panels contained nineteen and thirty-five measurements. The principal prediction comparisons reported here used scheduled noisy observation and the conventional panel; the phenotype–etiology analysis used the enriched panel. Broader sensitivity families retained their original panel and observation scopes. The shared Supplementary Methods provide the study-specific generating, observation, representation and statistical definitions (S1–S4).

For each scenario, four development cohorts of 1,024 individuals supported representation selection, and twenty independent confirmation cohorts of 5,115 individuals evaluated the locked procedures. Each confirmation cohort contained 3,069 training, 1,023 validation and 1,023 test individuals. The corresponding development partition was 614, 205 and 205. All records and future branches for an individual remained in the same partition. Repeated panels and observation conditions reused cohort identities and did not add independent population replications. The replication unit for complete inferential contrasts was twenty paired cohort-level values within a scenario and experimental context.

Measurements available through year 20 supported prediction of six conventional measurements at year 25. Continuation retained the specified loading inputs. The historical withdrawal branch reassigned inputs to a common low-load level of 0.05 from year 20; because initial inputs could be below that level, this assignment was not necessarily a reduction for every individual and domain. We refer to it as the low-load task while retaining the original branch identity in the analysis records. The two branches shared prior history and corresponding future innovations. These interventions were synthetic input rules, not clinical treatments or estimates of therapeutic effects.

### Information access and representations

A representation was constructed from the observed history and context available at the prediction cutoff. Predictive sufficiency would require the fuller history to add no information about the specified future outcome once the representation and common context were known (Dawid, 1979). Constructing a representation from history does not guarantee that condition: the construction can discard information. The present fitted-loss comparisons did not directly test equality of complete conditional distributions. Similarly, under squared-error loss, retaining a source from which a category can be calculated cannot worsen the best achievable conditional-mean prediction; a finite fitted procedure need not achieve that optimum. These distinctions separate information preservation from algorithm performance.

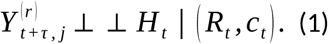

Current Measurements retained the latest eligible value in each channel. Current and Baseline Measurements additionally retained the observed baseline, but not intermediate measurement values. Complete Available Longitudinal Measurement History retained the eligible scheduled record and its support information. Current values could be older than the cutoff under the declared availability rules. Acquisition time and reporting time both had to occur on or before the cutoff; later observations could not repair earlier unsupported histories.

Training targets retained observed missingness, whereas principal held-out scoring used complete synthetic noisy ascertainment at year 25. Latent states and true hidden regimes were unavailable as predictor inputs.

Binary metabolic syndrome status and criterion count used the declared waist, triglyceride, HDL, pressure and glucose thresholds. Both pressure measurements contributed to a single criterion. Where criteria were missing, status could remain indeterminate and count retained lower and upper bounds; imputation for continuous prediction did not invent observed diagnostic information. These were operational classifications within the synthetic experiment. They did not establish a population-specific diagnostic standard beyond the stated conventions.

The mechanism-informed longitudinal representation Φ combined selected current values and historical descriptors. Its conventional-panel recipe retained all six current measurements and ten-year mean departures for systolic and diastolic pressure; baseline measurements, context and observation-support fields entered the prediction design separately. The wider-panel recipes contained sixteen and thirty-two selected values.

Development-stage coefficient information guided bounded selection, after which the recipe was fixed for confirmation. The descriptor rationale did not identify latent reserve, biological memory or causal mechanisms, and selection did not guarantee that Φ was optimal or sufficient.

Selected Measurements and Longitudinal History Descriptors drew named features from broader pools, retaining 48, 215 and 311 values across the three panels. The Outcome-Guided Longitudinal Embedding used a Ridge teacher fitted to permitted outcomes under both future tasks and a coefficient-based projection, with eight conventional-panel coordinates and twelve in each larger panel. Teacher fitting and cohort-specific projections respected the training and refitting boundaries. This was supervised linear compression, not unsupervised principal-components analysis. Neither the number of displayed coordinates nor the feature-family name identified a biological dimension.

Hard K-means memberships were fitted to specified source representations and supplied as one-hot labels. Standard cluster selection considered two to six groups using training-data geometry and minimum support. Separate sensitivities used predictive validation to select cluster number and diagonal-covariance Gaussian mixtures to compare hard assignments with soft membership vectors. Historical Cluster Membership and Transitions applied a frozen current-measurement map to eligible visits and retained membership, dwell, transition and coverage information. Baseline-only clusters used a separately fitted baseline map. Focused Glycaemic History used fasting-glucose history in the conventional panel and additional glycaemic channels only in panels containing them.

Input differences were retained explicitly. In the direct conventional-panel comparison, Φ received six raw baseline measurements absent from categorical history, while categorical history retained temporal memberships not supplied in that form to Φ. Both received their declared context and support fields. Thus, their comparison evaluated different longitudinal strategies. Comparisons between continuous measurements and clusters derived from those same measurements addressed aggregation more directly, although encoding, tuning and learner behaviour still differed. No result was interpreted as isolating continuity merely because one method’s displayed output was continuous.

### Predictive learning and inferential procedures

Primary prediction used Ridge regression with an unpenalised intercept. Validation selected among nine penalties from 10⁻⁴ to 10⁴ using equal-output, scale-normalised loss; the selected configuration was then refitted on training plus validation. Histogram-based gradient boosting supplied a bounded nonlinear sensitivity, using 200 iterations, learning rate 0.05 and six combinations of leaf count and L2 penalty. Different arms could select different permitted settings. Imputation, scaling, clustering and supervised projections were fitted only on the appropriate fitting population. Test outcomes could not select representations, revise projections or tune the prediction procedure.

For each cohort, task and channel, prediction loss was mean squared error divided by the squared standard deviation of the designated training-outcome population. The main aggregate weighted six channels equally. These dimensionless losses were not assay-error variances, clinical effect sizes or percentage errors. Channel-specific sensitivity endpoints were kept distinct from the six-channel average. Candidate-minus-comparator differences were formed within cohorts using common subjects, targets and normalisation; negative differences favoured the candidate. Complete nondegenerate comparisons used the mean paired difference, its sample standard deviation divided by the square root of twenty, a two-sided Student t test with nineteen degrees of freedom, and a pointwise 95% confidence interval. The t reference is exact for independent normal cohort contrasts and otherwise approximate.

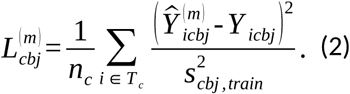

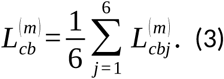

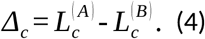

The joint primary Holm family remained 264 contrasts, including the historical 72-test subset without selecting whichever correction gave a favourable decision. The later Current Measurements-versus-own-clusters completion retained its separate twelve-test family and post-campaign status. Five sensitivity families retained 2,592, 1,944, 3,528, 324 and 7,920 hypotheses. Direct Φ-versus-categorical-history comparisons retained separate 72-test families for Ridge and boosting. The saved-comparator analysis, including formal learner interactions, retained its 468-test family. Dividing the results between two papers did not redefine these families. Pointwise intervals remained pointwise beside adjusted p-values, and family-specific error control did not provide simultaneous manuscript-wide control.

Differences in significance between learners were not treated as interaction tests. Formal interactions used the paired cohort contrast of the candidate-minus-comparator difference under boosting minus that difference under Ridge. Exact ties and incomplete support retained their original exceptional-case rules. Three original predictive tasks remained unavailable, leaving eighteen affected channel hypotheses without a verdict in the cluster-number sensitivity; their slots remained in the nominated multiplicity roster. These gaps were not replaced by favourable assumptions or later diagnostic results.

### Phenotype-bearing and recorded etiological information

The enriched-panel Ridge ablation compared three input designs under scheduled noisy observation. The exposure-and-demographic comparator contained the two observed etiological indicators together with baseline age and sex. Each indicator was correct when observed and independently missing with probability 0.20. The reduced phenotype-bearing design retained its registered values, baseline, support and demographic fields, while removing etiological indicators, controlled-input context, treatment, smoking and activity. The combined design added the two recorded indicators to that reduced phenotype block. Consequently, the comparator was not complete etiology and the phenotype arm was not the ordinary full Φ design.

All twenty cohorts and 1,023 test individuals per cohort contributed in each scenario and task. The release supplied thirty-six component losses and twenty-four paired contrasts using the six-channel standardised endpoint. These results were descriptive: no additional significance tests, confidence intervals, equivalence margin or causal mediation analysis was introduced. Adding or removing a predictor field changed what the learner observed, without removing that field’s generating influence from the synthetic physiology.

### Post hoc assessment of transition preservation

A separate mathematical analysis queried the original monthly current-state-sufficient law under noiseless conventional measurement and one fixed admissible context. It concerned constructed states at year 1, not sampled individuals at the year-20 prediction cutoff. Two pairs were selected before probability evaluation; each pair shared current binary status and criterion count. For a destination category, the category-transition probability was obtained by integrating the state-transition kernel over all next states mapped to that category. Strong lumpability required equality of every destination-category probability for every admissible pair in the same origin category under the same context and input rule (Kemeny and Snell, 1960; Geiger and Temmel, 2014).

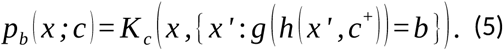

Deterministic integration retained shared Gaussian innovations, dependence between the triglyceride and HDL criteria, and the joint blood-pressure rule. Numerical convergence estimates qualified the calculated magnitudes; they were not confidence intervals or certified numerical enclosures. An independent monotone-coupling argument established strict inequality on a positive-probability open region. Supplementary Methods S6 specifies the law, category events, admissibility and proof. The analysis remained post hoc to the original programme. It neither changed the annual observation schedule nor established the Markov property of a category process under a particular initial population distribution.

## Results

### Categorisation and fitted prediction

Under scheduled noisy observation, the conventional panel and Ridge regression, Φ had lower six-channel loss than binary syndrome status and criterion count in all twelve scenario–task comparisons against each comparator; all rejected under Holm-264. In current-state-sufficient continuation, Φ minus binary status was −0.0156161 (pointwise 95% CI −0.0187030 to −0.0125293; adjusted p = 2.89 × 10⁻⁷). The corresponding count contrast was −0.0151545 (−0.0170971 to −0.0132120; adjusted p = 2.43 × 10⁻¹⁰). Table 2 and Figure 1 retain the information conditions and selected adverse and non-rejected comparisons.

**Figure 1.**
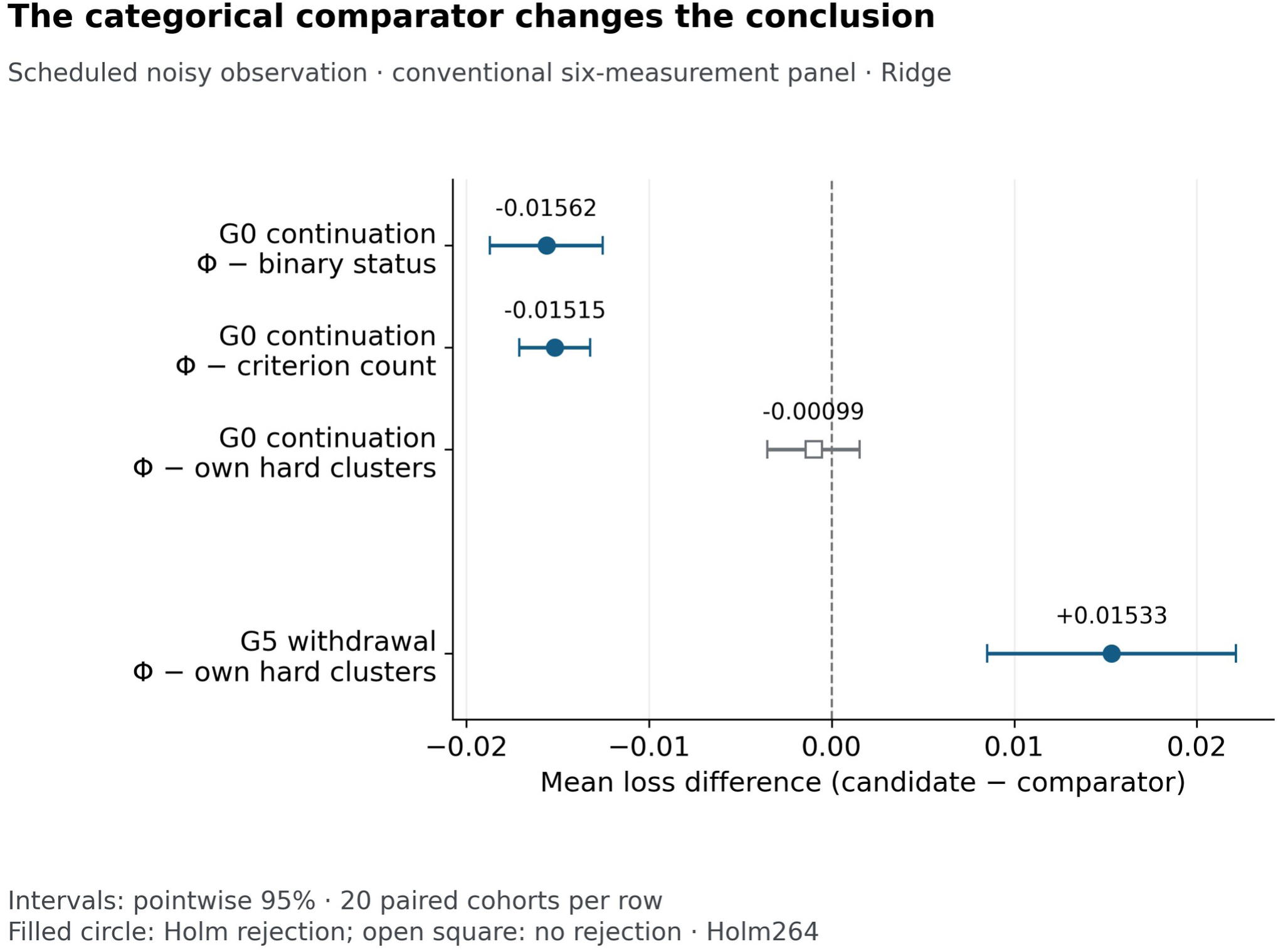
The predictive comparison depended on the source and comparator. Selected scheduled-noisy conventional-panel Ridge contrasts show lower Φ loss than syndrome status and count, a non-rejected own-cluster comparison in current-state-sufficient continuation, and lower cluster loss in nonlinear low-load prediction. Points are equal-cohort six-channel mean-loss differences; lines are pointwise 95% confidence intervals from twenty paired cohorts. Holm-264 decisions remain unchanged. Negative values favour Φ; the input information was not identical across all comparisons. Historical “withdrawal” labels, where shown, denote assignment to the specified low-load regime.

**Table 2.** Selected primary contrasts showing the importance of comparator and context.

| Scenario and task | Candidate minus comparator | Mean loss difference | Pointwise 95% CI | Holm-264 p |
| --- | --- | --- | --- | --- |
| Current-state-sufficient; continuation | $\Phi$ – binary MetS status | -0.0156161 | -0.0187030 to -0.0125293 | $2.89 \times 10^{-7}$ |
| Current-state-sufficient; continuation | $\Phi$ – MetS criterion count | -0.0151545 | -0.0170971 to -0.0132120 | $2.43 \times 10^{-10}$ |
| Current-state-sufficient; continuation | $\Phi$ – its own hard clusters | -0.0009868 | -0.0035087 to +0.0015351 | 1.000 |
| Nonlinear, regime-dependent; low-load | $\Phi$ – its own hard clusters | +0.0153295 | +0.0085174 to +0.0221415 | 0.01280 |
| Current-state-sufficient; continuation | $\Phi$ – Current and Baseline Measurements | -0.0000369 | -0.0008696 to +0.0007958 | 1.000 |
| Accumulated burden; continuation | $\Phi$ – Current Measurements | -0.0059490 | -0.0071074 to -0.0047907 | $2.30 \times 10^{-7}$ |
| Accumulated burden; continuation | $\Phi$ – Current and Baseline Measurements | +0.0003520 | -0.0004745 to +0.0011785 | 1.000 |

Scheduled noisy conventional-panel observation; Ridge; twenty paired cohorts; equal six-channel standardised loss. Negative differences favour the first named method. Confidence intervals are pointwise. The original Holm-264 family is unchanged. Input access was not identical in all comparisons. These selected rows do not replace the complete family.

Own-source comparisons were more mixed. None of Φ’s six continuation comparisons with its own clusters rejected under the joint primary correction. Low-load prediction favoured Φ in four scenarios, was non-rejected in adaptive-reference dynamics, and favoured its clusters in nonlinear dynamics. For the latter, Φ minus its clusters was +0.0153295 (+0.0085174 to +0.0221415; adjusted p = 0.01280). Selected descriptors outperformed their clusters in ten comparisons; both nonlinear comparisons were non-rejected. The learned embedding outperformed its clusters in eleven comparisons, with nonlinear low-load prediction non-rejected. These results did not establish a learner-independent ordering of information content.

The separate Current Measurements-versus-own-clusters completion was more consistently favourable to continuous inputs. All twelve mean contrasts were negative, with eleven rejecting under Holm-12; adaptive-reference continuation was non-rejected. Table 3 retains the complete family. Nonlinear continuation and low-load differences were −0.084967 and −0.094203, respectively. Because this comparison used clusters of the same current measurements, historical access was less prominent as an explanation than in Φ-versus-syndrome comparisons. Nevertheless, it remained an evaluation of fitted procedures, and the analysis was completed after related results were known.

**Table 3.** Current measurements compared with clusters of those same measurements.

| Biological scenario | Future task | Mean loss difference | Pointwise 95% CI | Holm-12 adjusted p |
| --- | --- | --- | --- | --- |
| Current-state-sufficient | Continuation | -0.005202 | -0.007825 to -0.002580 | 0.001083 |
| Current-state-sufficient | Low-load | -0.023799 | -0.027289 to -0.020309 | $1.31 \times 10^{-10}$ |
| Accumulated burden | Continuation | -0.004611 | -0.006588 to -0.002635 | 0.000413 |
| Accumulated burden | Low-load | -0.015475 | -0.018981 to -0.011970 | $1.30 \times 10^{-7}$ |
| Reserve depletion and recovery | Continuation | -0.004182 | -0.006077 to -0.002288 | 0.000560 |
| Reserve depletion and recovery | Low-load | -0.017791 | -0.020868 to -0.014714 | $2.02 \times 10^{-9}$ |
| Adaptive physiological reference | Continuation | -0.002021 | -0.004770 to +0.000728 | 0.1404 |
| Adaptive physiological reference | Low-load | -0.012153 | -0.016380 to -0.007926 | 0.0000433 |
| Persistent historical effects | Continuation | -0.007212 | -0.009433 to -0.004991 | 0.0000103 |
| Persistent historical effects | Low-load | -0.019180 | -0.022995 to -0.015365 | $1.84 \times 10^{-8}$ |
| Nonlinear, regime-dependent | Continuation | -0.084967 | -0.090532 to -0.079403 | $6.67 \times 10^{-17}$ |
| Nonlinear, regime-dependent | Low-load | -0.094203 | -0.103751 to -0.084655 | $1.96 \times 10^{-13}$ |

Complete twelve-comparison post-campaign family, scheduled noisy conventional-panel observation and Ridge. Differences are Current Measurements minus their own hard clusters. Twenty paired cohorts per comparison. Pointwise 95% intervals accompany the separate Holm-12 correction. Non-rejection is not equivalence.

Shared labels also contained variation in future measurements. Across the released continuation waist summaries, within-category variance ranged from 25.5084 to 65.7843 cm² for binary status and from 32.2868 to 60.6308 cm² for current-measurement clusters, with 2,960 finite records in each range. This variation included process and observation randomness; its existence did not establish that all of it was predictable from distinctions omitted by the category.

Historical comparisons also found latent change within unchanged categories. In accumulated-burden dynamics under scheduled noisy conventional-panel observation, material latent change from years 10 to 20 occurred among 27.18% of unchanged binary labels, 21.45% of unchanged criterion counts and 27.93% of unchanged current-measurement clusters. Conversely, among cases meeting the latent-change rule, 74.47%, 40.35% and 71.34% retained those respective labels. Material change meant D ≥ 0.5, where D was the root mean square of four burden changes standardised by their training-population year-20 standard deviations. These were equal-cohort means across twenty evaluable cohorts, not pooled individual proportions. The two sets use different denominators; the threshold was operational, not clinical. These findings describe concealed historical change, not a newly estimated effect of category silence on predictive loss.

### Historical information beyond current measurements

Φ outperformed Current Measurements in all twelve primary scenario–task comparisons under Holm-264, but none of its comparisons with Current and Baseline Measurements rejected. In accumulated-burden continuation, Φ minus current measurements was −0.0059490 (−0.0071074 to −0.0047907; adjusted p = 2.30 × 10⁻⁷), whereas Φ minus current-plus-baseline measurements was +0.0003520 (−0.0004745 to +0.0011785; adjusted p = 1). Figure 2 illustrates why the reference information set changes the conclusion. Non-rejection did not establish that baseline measurements contained every useful aspect of history.

**Figure 2.**
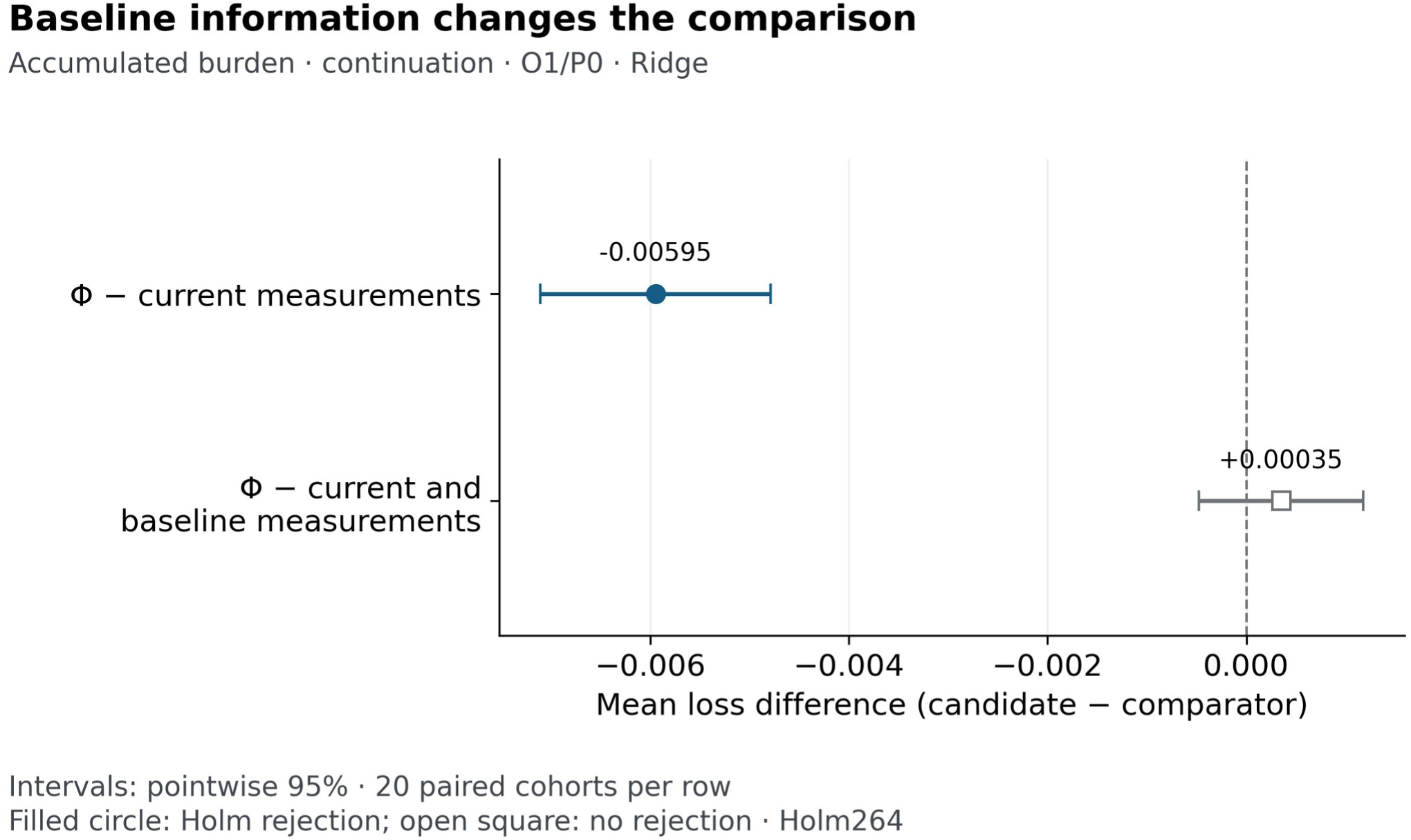
A benefit over current measurements did not establish a benefit over current plus baseline measurements. In accumulated-burden continuation, Φ improved six-channel standardised loss against current measurements, while its baseline-augmented comparison was non-rejected. Points and pointwise 95% intervals use twenty paired cohorts under scheduled noisy conventional-panel observation and Ridge. Holm-264 decisions are retained; non-rejection does not establish equivalence. Historical “withdrawal” labels, where shown, denote assignment to the specified low-load regime.

The correction scope also mattered. In current-state-sufficient low-load prediction, the Φ-minus-current-plus-baseline pointwise interval was entirely negative, from −0.0036253 to −0.0010763. Its joint Holm-264 p-value was 0.07376, whereas the retained historical Holm-72 value was 0.02002. The primary conclusion followed the joint family. This was one result viewed under two historical correction scopes, not two independent findings.

Selected descriptors and the learned embedding showed benefits over current-plus-baseline measurements in specified contexts. The embedding improved both tasks in current-state-sufficient and persistent-historical-effects dynamics, and continuation in adaptive-reference dynamics. Selected descriptors improved low-load prediction in the first four scenarios and both tasks in persistent-historical-effects dynamics. Neither construction established a universal advantage. Useful history in the current-state-sufficient scenario was compatible with its Markov generating state: earlier observations could inform an imperfectly measured present without requiring an additional active historical carrier.

Categorical history also retained useful temporal information. In nonlinear continuation fasting glucose, Historical Cluster Membership and Transitions outperformed current membership by −0.0955080 (−0.1045650 to −0.0864506; Holm-3,528 p = 1.75 × 10⁻¹¹) and baseline membership by −0.0879249 (−0.0921687 to −0.0836811; adjusted p = 6.37 × 10⁻¹⁷). These were channel-specific contrasts, not six-channel averages. The separately fitted baseline map also differed from the frozen map used across history. Corresponding current-state-sufficient continuation comparisons did not establish the same benefit, so the nonlinear example did not support a universal history advantage.

### Recorded etiological information and phenotype-bearing inputs

The combined design had lower saved loss than the limited exposure-and-demographic comparator in every scenario–task context. In current-state-sufficient continuation, mean losses were 0.690890 for recorded etiology with demographics, 0.231081 for reduced phenotype-bearing information and 0.228898 for the combination.

These were equal-cohort means of the six-channel standardised endpoint. They did not identify which individual measurement, historical descriptor, baseline value or support field supplied the difference.

Adding recorded etiology to the reduced phenotype produced smaller, context-dependent changes. All six continuation contexts improved, as did three low-load contexts. The combined design had higher loss in current-state-sufficient, accumulated-burden and nonlinear low-load prediction, with paired increments +0.000155601, +0.000274448 and +0.000309893, respectively. Table 4 retains all component means and incremental contrasts, and Figure 3 displays the twelve increments. These descriptive differences do not establish significance, equivalence, etiological redundancy or negative information.

**Figure 3.**
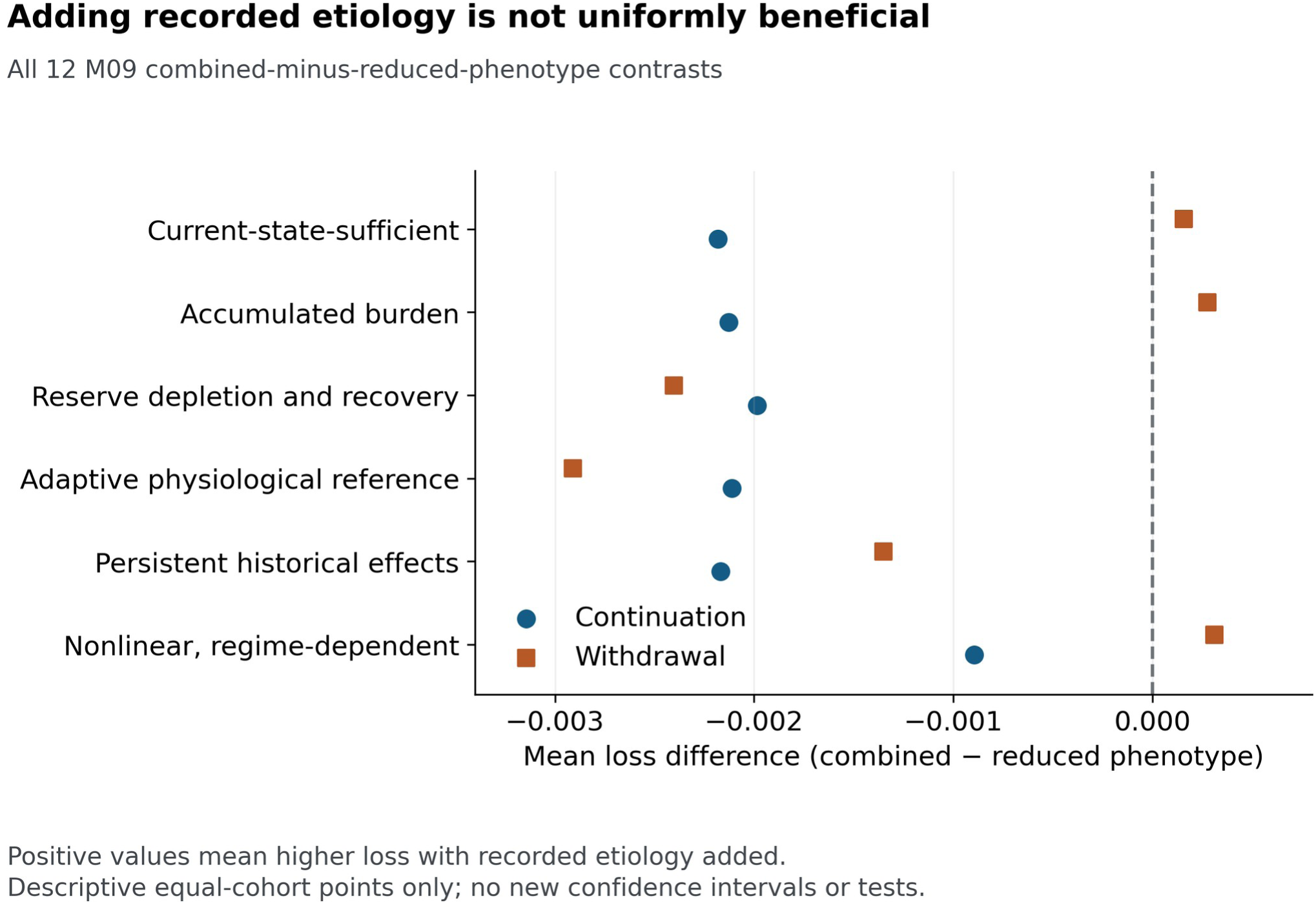
The incremental contribution of recorded etiology depended on scenario and future task. Adding two recorded exposure indicators to the reduced phenotype-bearing design lowered saved loss in all continuation contexts and increased it in three low-load contexts. Points reproduce all twelve equal-cohort incremental contrasts from the scheduled noisy enriched-panel Ridge analysis. Negative values favour the combined design. These descriptive estimates have no added confidence intervals or significance tests. M09 is the original identifier for the phenotype–etiology analysis. Historical “withdrawal” labels, where shown, denote assignment to the specified low-load regime.

**Table 4.** Complete phenotype and recorded-etiology component losses and incremental contrasts.

| Scenario | Task | Etiology and demographics | Reduced phenotype | Combined | Combined minus phenotype |
| --- | --- | --- | --- | --- | --- |
| Current-state-sufficient | Continuation | 0.690890 | 0.231081 | 0.228898 | -0.002182870 |
| Current-state-sufficient | Low-load | 0.695193 | 0.531714 | 0.531870 | +0.000155601 |
| Accumulated burden | Continuation | 0.711102 | 0.187436 | 0.185307 | -0.002129127 |
| Accumulated burden | Low-load | 0.689084 | 0.387215 | 0.387489 | +0.000274448 |
| Reserve depletion and recovery | Continuation | 0.681406 | 0.199765 | 0.197778 | -0.001986047 |
| Reserve depletion and recovery | Low-load | 0.664425 | 0.451384 | 0.448982 | -0.002402358 |
| Adaptive physiological reference | Continuation | 0.696012 | 0.178843 | 0.176731 | -0.002112371 |
| Adaptive physiological reference | Low-load | 0.676522 | 0.379733 | 0.376824 | -0.002908829 |
| Persistent historical effects | Continuation | 0.714787 | 0.175053 | 0.172886 | -0.002167612 |
| Persistent historical effects | Low-load | 0.691689 | 0.368122 | 0.366770 | -0.001352115 |
| Nonlinear, regime-dependent | Continuation | 0.790067 | 0.158766 | 0.157871 | -0.000894837 |
| Nonlinear, regime-dependent | Low-load | 0.769392 | 0.262486 | 0.262795 | +0.000309893 |

Scheduled noisy enriched-panel Ridge analysis, twenty cohorts and 1,023 held-out individuals per cohort. Component means are displayed to six decimals. The last column contains saved paired-cohort contrasts, not new subtraction of the rounded means. All are descriptive; no confidence intervals or significance tests were added. The phenotype design includes baseline, support and demographic information; the etiological comparator contains two observed indicators with age and sex.

The comparison was asymmetrical by design: two incompletely observed exposure indicators were compared with a larger block containing measured phenotype, baseline and support information. Their limited incremental predictive value did not rank the causal or therapeutic importance of the underlying influences. Because exposure remained part of the generating process after its indicator was removed from a predictor, the ablation distinguished information access from intervention on physiology.

### Stronger alternatives and learner dependence

Strengthening categorical alternatives changed particular comparisons without producing a general winner. In nonlinear continuation fasting glucose, soft minus hard current-measurement mixture membership had a loss difference of −0.0144546 (−0.0157444 to −0.0131648; Holm-1,944 p = 3.35 × 10⁻¹²). Predictive rather than geometric selection of current-measurement cluster number yielded −0.0488887 (−0.0566408 to −0.0411367; Holm-324 p = 1.64 × 10⁻⁸). The corresponding learned-embedding-source cluster-number comparison was non-rejected despite a negative pointwise interval. Table 5 reports selected contrasts with their distinct correction families.

**Table 5.** Selected stronger-comparator and learner sensitivities.

| Candidate – comparator | Learner | Mean loss difference | Pointwise 95% CI | Holm-adjusted p |
| --- | --- | --- | --- | --- |
| $\Phi$ – its own hard clusters | Ridge | +0.0013934 | -0.0021392 to +0.0049260 | 1.000 |
| $\Phi$ – its own hard clusters | Boosting | +0.0107904 | +0.0073989 to +0.0141819 | 0.003309 |
| Soft – hard current-measurement mixture membership | Ridge | -0.0144546 | -0.0157444 to -0.0131648 | $3.35 \times 10^{-12}$ |
| Outcome-selected – standard current-measurement cluster number | Ridge | -0.0488887 | -0.0566408 to -0.0411367 | $1.64 \times 10^{-8}$ |
| Focused glycaemic history – $\Phi$ | Ridge | -0.0004247 | -0.0027247 to +0.0018753 | 1.000 |
| Focused glycaemic history – $\Phi$ | Boosting | -0.0080394 | -0.0102699 to -0.0058088 | 0.001828 |

Nonlinear continuation fasting glucose; scheduled noisy conventional-panel observation; twenty paired cohorts. This is a channel-specific endpoint. $\Phi$ versus clusters uses Holm-2,592; soft versus hard membership uses Holm-1,944; cluster-number selection uses Holm-324; focused glycaemic history uses Holm-7,920. Families are not pooled.

Learner choice also changed the comparison with a focused physiological history. In the same nonlinear continuation glucose context, Focused Glycaemic History minus Φ was −0.0004247 under Ridge, with an interval spanning zero and adjusted p = 1. Under boosting it was −0.0080394 (−0.0102699 to −0.0058088; Holm-7,920 p = 0.001828). Conventional-panel focused history used glucose observations only; it did not borrow insulin, glycated haemoglobin or two-hour glucose from larger panels. Under boosting, Φ also had higher glucose loss than its own clusters in this context. Differences between significance labels alone were not treated as formal learner interactions.

The direct Φ-versus-categorical-history analysis contained 72 complete channel contrasts per learner. Ridge yielded 50 rejections favouring Φ, eleven favouring categorical history and eleven non-rejections. Boosting yielded 56, thirteen and three, respectively. Categorical history retained systematic favourable contexts: under Ridge, waist circumference in both tasks in the first five scenarios and in nonlinear low-load prediction; under boosting, continuation waist in all scenarios and continuation HDL in the first five. Counts of decisions reused cohorts and outcomes and were not independent votes or probabilities of superiority.

Eight direct contrasts changed sign across learners, including four with opposite-direction rejections. For nonlinear low-load waist, Φ minus categorical history changed from +0.0067992 under Ridge to −0.0058421 under boosting (Figure 4). These reversals concerned differently constructed longitudinal inputs, because Φ had raw baseline measurements absent from categorical history. The largest favourable direct Φ contrast was current-state-sufficient low-load diastolic pressure under both learners: −0.1019952 for Ridge and −0.1047035 for boosting. Conversely, the largest positive Ridge contrast, nonlinear low-load triglycerides at +0.0185525, remained non-rejected after Holm-72 correction.

**Figure 4.**
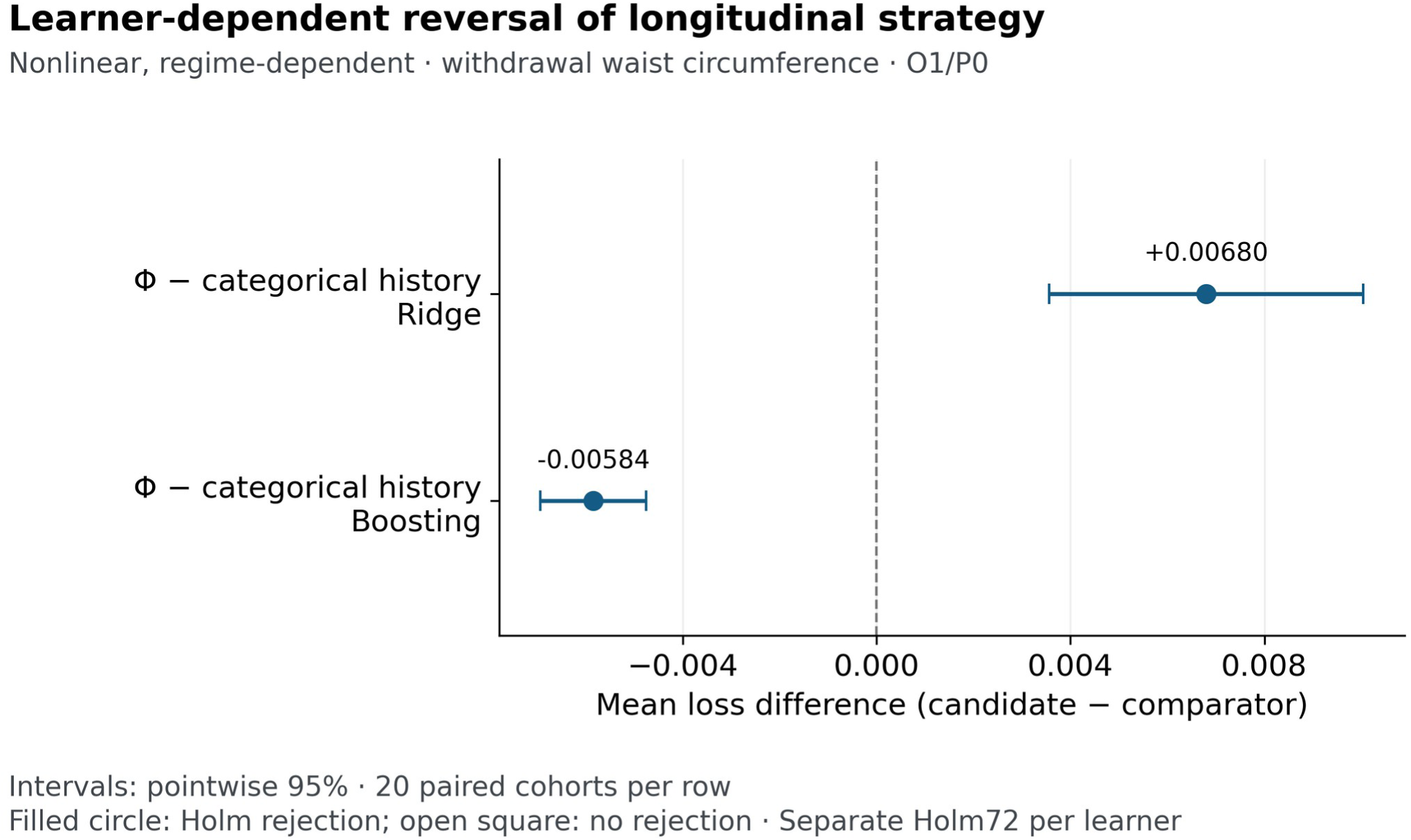
The ordering of two longitudinal strategies reversed between learners. For nonlinear low-load waist circumference, Ridge favoured categorical history and boosting favoured Φ. Points and pointwise 95% intervals use twenty paired cohorts under scheduled noisy conventional-panel observation and separate Holm-72 families. Φ and categorical history had different baseline and temporal information. This descriptive reversal is not a separately tested interaction. Negative values favour Φ. Historical “withdrawal” labels, where shown, denote assignment to the specified low-load regime.

Formal interaction evidence came from a different family. In nonlinear low-load prediction, the six-channel Φ-minus-own-cluster contrast was −0.0062805 under boosting and +0.0153295 under Ridge. Their cohort-paired interaction was −0.0216100 (−0.0292503 to −0.0139697; Holm-468 p = 0.002310). Current-state-sufficient continuation Current Measurements-minus-own-clusters had a non-rejected interaction of +0.0005812. These tests established specific learner dependencies and were not transferred to the channel-specific direct comparisons, where interaction tests had not been performed.

### Categories did not preserve the examined monthly transition law

All four constructed current-state-sufficient probes were syndrome-negative with criterion count two. Within each pair, the second state had greater burden in all four domains. The pairs shared the same female-threshold context, no qualifying treatment flags and a fixed input rule. They were mathematical queries within the support of the year-1 process, not four selected cohort participants. Their current profiles lay strictly inside their respective categories, and sufficiently small neighborhoods preserved those categories.

For the first pair, next-month syndrome-positive probabilities were 0.10157085 and 0.69537270, a difference of 0.59380185, or approximately 59.38 percentage points. The second pair gave 0.09907892 and 0.69359219.

Table 6 and Figure 5 retain these model-kernel probabilities. The corresponding numerical difference-error estimates were approximately 1.10 × 10⁻¹⁰ and 2.11 × 10⁻¹⁰, interpreted as convergence estimates rather than certified bounds. An independent proof established strict inequality: under shared innovations, the ordered second state could not have fewer future positive criteria, and a nonempty open region gave the first exactly two criteria and the second at least three.

**Figure 5.**
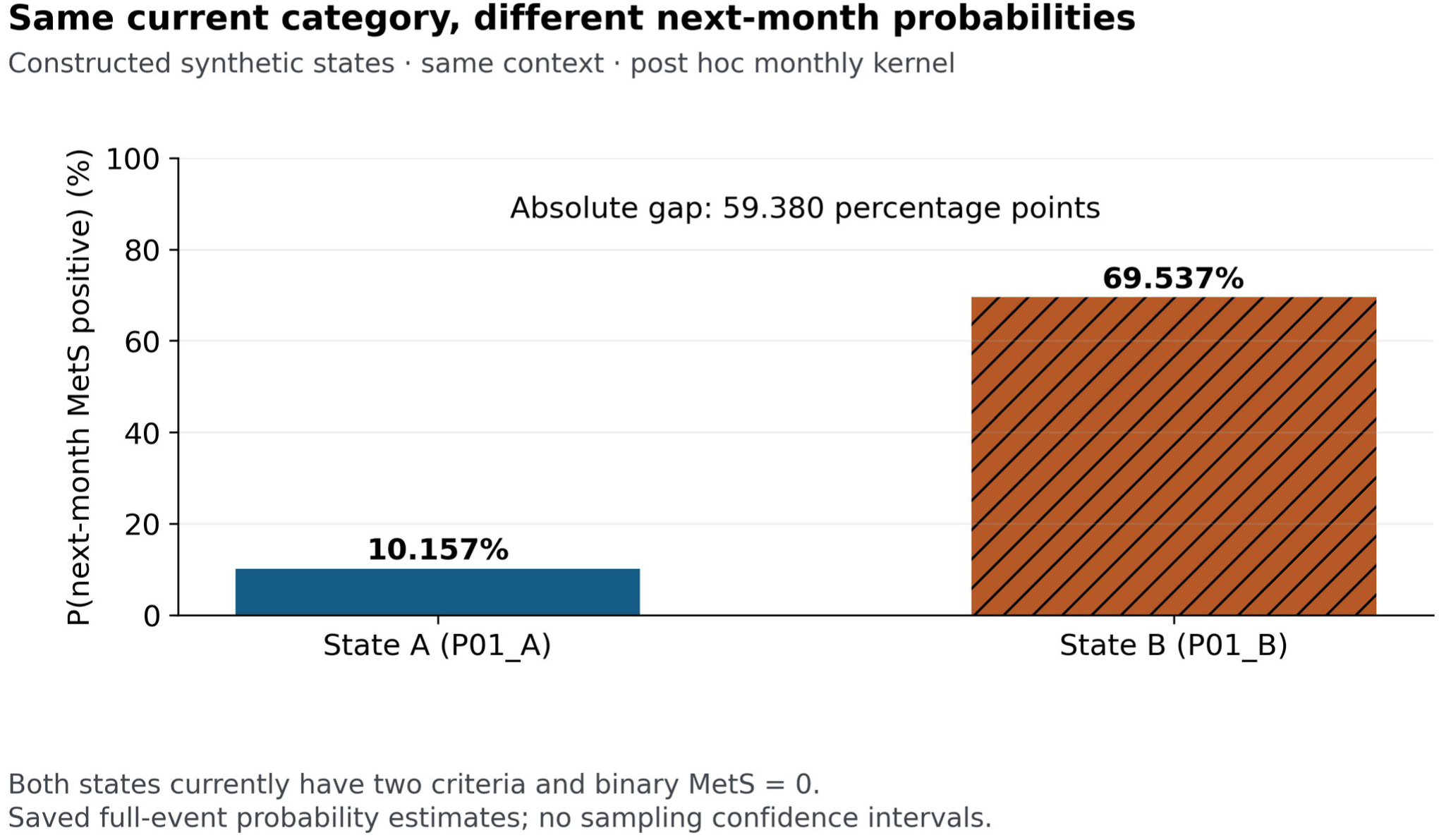
Equal current status and count did not determine equal transition probabilities. Two constructed states, both syndrome-negative with count two, had next-month positive-category probabilities of 10.16% and 69.54% in the same fixed context under the current-state-sufficient law. Numerical integration quantified the difference; an independent analytic argument proved strict inequality. These are synthetic kernel probabilities, not patient risks or population-average effects. The monthly analysis is distinct from the five-year predictive experiment.

**Table 6.** Shared present categories with unequal next-month category probabilities.

| Constructed state | Current glucose mg/dL | Current criterion count | Current syndrome status | Next-month positive probability |
| --- | --- | --- | --- | --- |
| Pair 1 A | 99.0 | 2 | Negative | 0.10157085 |
| Pair 1 B | 99.5 | 2 | Negative | 0.69537270 |
| Pair 2 A | 99.0 | 2 | Negative | 0.09907892 |
| Pair 2 B | 99.5 | 2 | Negative | 0.69359219 |

Constructed admissible states under the current-state-sufficient monthly law. All use the same female-threshold context and no qualifying treatment. Probabilities integrate the biological innovation law with noiseless destination measurements; they are not observed human risks. Full states and classification events are given in Supplementary Methods S6.

The binary transition probabilities therefore differed within a shared current binary category. The future count distributions also differed, because their probabilities summed over counts three to five were unequal despite equal current counts. Both partitions failed strong lumpability for the specified one-month law and context. The claim did not depend on interpreting the large selected numerical gap as typical, and the second pair did not constitute population replication.

A later marginal calculation explained much of the first pair’s numerical contrast. Current glucose values of 99.0 and 99.5 mg/dL became deterministic pre-innovation values of approximately 99.64964 and 100.14201 mg/dL. Their glucose-threshold crossing probabilities were approximately 0.10157085 and 0.69535601, close to the joint syndrome probabilities. All four starting burdens differed and other domains influenced the glucose target, so this calculation did not isolate a causal glucose contribution. The intermediate values were not necessarily expectations after stochastic transformation. The result did not establish a physiological bifurcation, clinical consequences of a small human glucose difference, annual non-lumpability or five-year predictive superiority.

## Discussion

### Representation must be evaluated for a specified future

The principal finding is conditional rather than a ranking of representation families. Conventional labels could conceal distinctions relevant to future metabolic measurements, and Φ improved prediction relative to those labels under the principal conditions. Yet Φ-derived clusters outperformed Φ in nonlinear low-load prediction, stronger categorical constructions recovered useful information, and the result depended on the learner. Continuous values, historical descriptors and categories should therefore be assessed by what information they receive and how their intended prediction procedure uses it.

The distinction between information and fitted performance explains why apparently opposing findings can coexist. A deterministic category cannot contain more information than the source from which it is calculated under common context. It can nevertheless be easier for a particular finite learner to use. Encoding, regularisation, restricted functional form and estimation variability are plausible contributors to the observed reversals, but their individual contributions were not identified here. Our findings extend the established concerns about categorisation into controlled longitudinal settings; they do not establish that the practical value of clinical categories is exhausted by their predictive loss.

The human literature gives this question substantive motivation. Franco’s within-syndrome heterogeneity and the value of continuous features in comparisons with diabetes subgroups concern clinical populations and outcomes (Franco et al., 2009; Dennis et al., 2019). The present outcomes were synthetic future measurements, not cardiovascular events or human disease progression. The connection is the question of what a common label leaves unresolved. A model with known generating laws permits that question to be examined under controlled assumptions, but it cannot substitute for empirical calibration or external evaluation.

### History and recorded causes answer different information questions

History was useful only relative to what the comparator already knew. Φ’s advantage over current measurements did not establish an advantage over current plus baseline measurements under the joint primary correction. More extensive longitudinal constructions and categorical histories supplied additional benefits in specified settings, but no benefit identified a unique biological memory mechanism. In the current-state-sufficient scenario, historical observations could improve estimation of an imperfectly observed present. They could also inform persistent subject–channel measurement effects, because the main targets were noisy future measurements. The experiment did not apportion gains between those explanations.

The distinction complements human longitudinal research. Scuteri and Tabák established temporal structure around metabolic disease development, while Breeze fitted correlated metabolic trajectories to human observations (Scuteri et al., 2009; Tabák et al., 2009; Breeze et al., 2016). A fixed-cutoff forecast requires that all predictor information already be available at the forecasting time. The present comparisons enforced that boundary, but they did not reproduce those studies’ clinical endpoints or establish which synthetic carrier corresponds to a human mechanism. Predictive success and biological explanation remain different aims (Shmueli, 2010).

The phenotype–etiology ablation reinforced the need to specify the whole input block. Recorded indicators omitted much of the information contained in the measurements, baseline and support fields, so their weaker prediction did not make causes unimportant. Small or adverse incremental differences could reflect the fitted procedure and available evidence without implying causal redundancy. Future empirical comparisons should balance access deliberately where the scientific question concerns transformation, while separately evaluating broader information strategies where that is the intended clinical comparison.

### Transition preservation and practical prediction

The lumpability result addresses a property of the partition and transition law. Its proof shows that some distinctions removed by the categories matter to their own next-month evolution, even when present burden is sufficient for the underlying dynamics and noiseless measurements retain it. This is stronger than observing multiple values inside a category, because the next-category probabilities were shown to differ. It is also narrower than a claim that categories are universally poor predictors. The five-year loss experiments used different outcomes, observation conditions and learners.

Empirically fitted category-transition models can remain useful despite the lack of exact preservation for a particular synthetic law. Lin and colleagues estimated transitions among metabolic categories and cardiovascular outcomes in human screening data (Lin et al., 2021). Our counterexample does not invalidate those associations or determine the Markov property of their observed process. Failure of strong lumpability is a failure of a universal equality across admissible states; it does not establish non-Markovianity under every initial distribution. Neither the proof nor its selected probability gap determines clinical importance, typical prevalence or the best replacement representation.

### Scope limitations and subsequent evaluation

The main limitation is the stipulated generating system. Transition coefficients, distributions and observation rules were not fitted to joint human metabolic trajectories. Known loading and future regimes supplied information that may be unavailable in clinical records. Repeated cohorts quantified variation under those laws, not uncertainty about whether the laws describe humans. The joint observation-and-treatment stress condition also changed physiology, so it could not isolate measurement noise. Equal channel weights and training-variance normalisation defined experimental losses rather than clinical utilities.

The analysis evaluated bounded constructions rather than an exhaustive search. Development used Ridge-based information to select Φ and the learned embedding; evaluating those constructions with boosting did not redesign each representation for that learner. Stronger comparator sensitivities and the mathematical investigation retained their later chronology. Three unavailable original predictions and the affected no-verdict comparisons remain part of the evidence. Exact historical mixture initialisation and dependency provenance were incomplete, limiting claims of exact retraining. Numerical agreement with an implementation would not itself establish biological validity. The supplied method definitions permit inspection of the experiment, but full computational reproduction additionally requires executable and numerical assets described in Supplementary Methods S7.

The companion paper evaluates predictive adequacy and representation movement using the same parent programme. Those properties should not be inferred from the present favourable comparisons: the evaluated Φ common recipes did not receive a complete predictive-adequacy certificate in that analysis. Measurement requirements also extend beyond displayed coordinates to baseline, support and source observations. We therefore make no claim of sufficient physiology or reduced clinical acquisition on the basis of this paper’s prediction results. The two papers address complementary endpoints and reuse cohorts rather than constituting independent validation studies.

Subsequent work should define a population, measurement process, prediction cutoff, horizon and future treatment context before comparing representations. Deliberately matched input comparisons would separate transformation from additional information access, and independent data would be needed to evaluate procedures designed after inspecting these results. Probabilistic extensions should assess calibration as well as predictive sharpness (Gneiting et al., 2007). Questions about treatment require distinct causal estimands and cannot be answered by predictor ablation alone (van Geloven et al., 2020). The present contribution is a controlled experiment showing how such distinctions can be examined while retaining favourable, adverse and unsupported outcomes.

## Conclusions

Representations of metabolic observations preserved different distinctions under the examined synthetic dynamics. Categories could conceal future-relevant information without being uniformly inferior for fitted prediction, while longitudinal information helped only relative to the comparator’s existing access. Recorded etiological indicators and phenotype-bearing inputs answered different predictive questions. A separate monthly-kernel counterexample established failure of strong lumpability for conventional status and count under one specified law. Together, these results support evaluating metabolic representations against defined information sources, tasks and learners. They do not establish a sufficient human physiological state, universal superiority of Φ, replacement of metabolic syndrome in clinical care, or therapeutic benefit.

## Declarations

### Relationship to the companion article

This article and the companion manuscript “Predictive adequacy and measurement dependence of metabolic representations in controlled synthetic systems” report distinct questions from the same synthetic study programme. They share generating and observation methods and overlapping simulated cohorts; they are not independent replications. The shared Supplementary Methods document specifies the bounded study design.

### Ethics and consent

The study used computationally generated synthetic cohorts. It involved no recruitment of or interaction with human participants, no patient or participant records, no identifiable personal information or biological specimens, and no animals. No ethics committee approval or formal exemption was obtained. Informed consent was not applicable.

### Data and code availability

The article and shared Supplementary Methods disclose the bounded synthetic model, observation mappings, locked Φ coordinates, representation and fitting rules, and the displayed result tables. The broader METLAS framework, confidential calibration resources and production implementation are outside the reported study and are not distributed.

The accompanying materials do not include the executable study implementation, complete selected-feature and embedding-input rosters, full-precision constructed-query records, complete machine-readable result families or paired-cohort loss arrays. Their absence limits exact computational reproduction. Requests for study-specific supporting materials may be directed to the corresponding authors and will be considered subject to ownership and release permissions; access is not guaranteed. Supplementary Methods S7 describes the materials provided and the remaining reproducibility limitations.

## Author contributions

Dr Mwangi Njenga: Conceptualization, Methodology, Formal analysis, Investigation, Software, Writing, Original draft

Dr Waweru Peter: Conceptualization, Supervision, Writing-Review and Editing, Project administration

## Use of AI assistance

AI-assisted tools were used in computational implementation and analysis, and in the organisation, drafting and editing of the manuscripts. The authors are responsible for the scientific content, interpretation and final text.

## Supporting information

Shared study methods for the METLAS companion papers

