## Supplementary material for "Predictive consequences of alternative representations of metabolic measurements: A controlled synthetic study of categories, longitudinal information and transition preservation": Shared study methods for the METLAS companion papers

Dr Mwangi Njenga and Dr Waweru Peter
University of Nairobi, Nairobi, Kenya

Companion scope: Paper 1 examines categorisation, historical and recorded etiological information, prediction and monthly transition preservation. Paper 2 examines predictive adequacy, information requirements and representation movement. The two papers use shared simulations and are not independent validations.

This supplement describes the bounded synthetic study shared by the two companion papers. It specifies the generating assumptions, observation laws, retained representations, fitting and evaluation procedures, movement analyses and post hoc mathematical counterexample. Numerical parameters are experimental design assumptions; they were not fitted to human cohort joint distributions. A representation is a function of permitted observations and context, distinct from latent physiological state and from the fitted prediction model. The two papers reuse parts of one simulation programme and do not constitute independent replications.

### S1. Synthetic physiological generator

#### S1.1 State, cohorts and information boundaries

**Table S1. Latent state variables and domain order.**

| state | role |
|---|---|
| X_d | Current burden in domain d; domain order a (adiposity), g (glycaemic), l (lipoprotein), h (haemodynamic). |
| Q_d | Reserve; G2 burden target and restoration-rate carrier. |
| O_d | Adapted reference; G3 active target displacement. |
| M_d | Retained alteration; G4 active target displacement. |
| C_d | Cumulative exposure; G1 active target displacement. |

Each of six scenarios (G0–G5) had four development cohorts of 1,024 individuals and twenty independent confirmation cohorts of 5,115. Training/validation/test sizes were 614/205/205 in development and 3,069/1,023/1,023 in confirmation. One identity permutation per cohort was shared across eligible comparisons; an individual's records and branches stayed together. Training fitted parameters, validation selected permitted configurations, and final refitting used training plus validation with configuration fixed. Test outcomes were reserved for evaluation.

Baseline age was Uniform(18,30) years; the conventional female/male threshold category was Bernoulli(0.5), without a sex or gender generalizability claim. Follow-up was 35 years in monthly steps. Prediction used year-20 history for year-25 outcomes. The 120 confirmation scenario–cohort combinations contained 122,760 distinct held-out cohort–individual identities; repeated panels, conditions, channels, branches and replicas did not increase that count.

Predictors received available observations and permitted context; future outcomes were restricted to fitting or evaluation. Hidden states were diagnostic-only, and true etiological and regime labels were excluded. Recorded E1/E2 indicators were correct when available, each independently missing with probability 0.20. Missing exposure evidence was not converted into a diagnosis or probability. Compared arms shared public inputs and their declared future schedules.

#### S1.2 Initial states and controlled inputs

For domain $d$, $Z_d = \sqrt{0.3}\, z_0 + \sqrt{0.7}\, z_d$, with independent standard-normal components. Here $Z_d$ is the initialisation draw, distinct from the full physiological state vector. Initial burden is $X_d = sigmoid(-1.4 + 0.55\, Z_d)$; reserve $Q_d$ is $0.35 + 0.60\, Beta(4, 2)$; adapted reference $O_d$ is $0.15 + 0.25\, Beta(2, 4)$; retained alteration $M_d$ is $0.02 + 0.10\, Beta(1, 4)$; and cumulative exposure $C_d$ begins at zero. Beta draws are independent across subjects/domains unless a shared component is explicitly stated. The current-state-sufficient scenario fixes $Q = 0.75, O = 0.20, M = C = 0$. The nonlinear scenario draws hidden regime $K \in \{0, 1, 2\}$ uniformly and initializes $X_d = clip(c_K + 0.05\, Z_d, 0.02, 0.95)$, with centres $c_K \in \{0.12, 0.40, 0.68\}$.

The named common/domain Gaussian initialization draws also enter the input equation. Beta draws are independent across individuals/domains and from Gaussian draws unless explicitly shared. The independent noncausal covariate is LPA_i=exp[log(15)+0.7Z], Z standard normal. Here sigmoid(v)=1/(1+exp(−v)), logit(v)=log[v/(1−v)], and clip(v,a,b) restricts v to [a,b].

E1 and E2 are Bernoulli(0.20) and Bernoulli(0.15), smoking is Bernoulli(0.20), and activity is Uniform(0,1). The public controlled input is

$$U_d = clip\left(0.25 + 0.10\, z_0 + 0.10\, z_d + 0.15\mathrm{E1} + 0.10\mathrm{E2} + 0.08\, smoking - 0.08\, activity\, , 0.02\, , 0.80\right).$$

E1 denotes sustained-load context and E2 adverse drug/exposure context. Smoking and activity are held constant in this implementation. At year 20, the paired continuation branch retains its input and the low-load branch sets every input component to 0.05. The two branches share pre-year-20 history and future process innovations. The historical identifier "withdrawal" denotes this reassignment, not removal to zero: because original inputs can be as low as 0.02, the operation is not a decrease for every individual. The absolute and residual outcomes are interpreted using this precise intervention.

##### S1.3 Monthly transitions and scenario coefficients

Define $relax\left(v\, , a\, , k\, , \Delta t\right) = a + \left(v - a\right) e^{-k\Delta t}$ for initial value $v$, target $a$ and rate $k$, with $\Delta t = 1/12$ year. All updates use old states simultaneously. The historical carriers follow

$$C^{+} = 1 - \left(1 - C\right) e^{-X\Delta t/15},$$

$$Q^{+} = relax\left(Q\, , \frac{0.10}{0.10 + 0.18\, X + 0.02\mathrm{E2}}\, , 0.10 + 0.18\, X + 0.02\mathrm{E2}\, , \Delta t\right),$$

$$O^{+} = relax\left(O\, , 0.15 + 0.50\, X\, , 1/6\, , \Delta t\right),$$

$$M^{+} = relax\left(M\, , \frac{0.06\, X^2}{0.06\, X^2 + 0.01}\, , 0.06\, X^2 + 0.01\, , \Delta t\right).$$

For G0–G4, the target is the clipped sum of 0.05, $0.65\, U_d$, $0.08\, \overline{X}_{-d}$, $0.04\, clip\left(\left(age - 18\right)/50\, , 0\, , 1\right)$, and the active terms $w_C C_d + w_O\left(O_d - 0.20\right) + w_M M_d + w_Q\left(1 - Q_d\right)$, clipped to [0.01,0.95]. Scenario weights are G0 all zero; G1 $w_C = 0.25$; G2 $w_Q = 0.16$; G3 $w_O = 0.50$; G4 $w_M = 0.45$. Base relaxation rates are [0.18,0.22,0.25,0.30] per year, multiplied by $0.30 + 1.4\, Q_d$ only in G2. After relaxation, the logit receives innovation $0.06\sqrt{\Delta t}\left(\sqrt{0.3}\, \varepsilon_0 + \sqrt{0.7}\, \varepsilon_d\right)$, then the sigmoid restores a bounded state. Here $\overline{X}_{-d}$ is the mean burden in the other domains. Inactive carriers can be generated as diagnostics without being causal inputs to burden.

For G5, write the same bounded age term as $A = clip\left(\left(age - 18\right)/50\, , 0\, , 1\right)$ and define the following algebraic component:

$$B_d^{(5)} = 0.04 + b_K + 0.55\tanh\left(1.8\, U_d\right) + 0.20\, 1\left(\overline{X} > 0.45\right).$$

The replacement burden target is

$$T_d^{(5)} = clip\left(B_d^{(5)} + 0.10\sin\left(2\pi\, X_{d+1}\right) + 0.04\, A\, , 0.01\, , 0.95\right).$$

Here $b_K \in \left\{0\, , 0.10\, , 0.20\right\}$, $\overline{X}$ averages all four burdens, cyclic domain indexing is used, and rates are [0.12,0.30,0.65] according to regime. $B_d^{(5)}$ groups terms for notation; it is not an additional state variable. The same bounded stochastic innovation rule applies. This deliberately nonlinear scenario is not derived from engineered representation values.

Domain order is adiposity, glycaemic, lipoprotein and haemodynamic. Current age is baseline age plus elapsed time; the mean of other burdens excludes the updated domain, while the G5 interaction uses the next domain cyclically. The common and domain-specific Gaussian innovations have independent standard-normal components and are independent across months. All carrier and burden updates use old states and beginningof-month age simultaneously. No carrier is updated first to give its new value an artificial within-step effect on burden.

The scenarios respectively activate no historical carrier (G0), accumulated exposure (G1), reserve (G2), adapted reference (G3), retained alteration (G4), or the nonlinear mixture override (G5). Inactive carriers may supply observation diagnostics outside G0 without causing burden. G5 deliberately allows categorical success.

##### S1.4 Treatment-policy stress condition

O0, O1 and O2 keep all treatment flags false. O3 adds a simplified simulated policy: at annual checks before year 20, an inactive blood-pressure, triglyceride or glucose flag begins with probability 0.25 when the corresponding noiseless mapped SBP, TG or fasting-glucose value reaches 140, 200 or 126 in its native units. Flags persist. The next monthly update subtracts 0.15 times the appropriate active flag from U_h, U_l or U_g, with a zero lower bound. No low-HDL-specific treatment flag is introduced. After year 20, flags remain fixed and known in both branches; all compared arms receive the same flags.

The stress policy uses latent mapped measurements, whereas comparator criteria use available recorded observations. Learners do not receive those latent values. O3 changes both observation and physiology, so it cannot isolate measurement error or estimate clinical treatment effects.

### S2. Observation model and measurement dictionary

#### S2.1 Schedules, noise, missingness and availability

**Table S2. Scheduled measurement years by observation condition. Channels 0–18 comprise P0/P1; channels 19–34 are the additional P2 measurements.**

| Condition | Core channels 0–18 | Enriched channels 19–34 |
|---|---|---|
| O0 | Annual0–35 | Annual0–35 |
| O1 | 0,2,5,7,10,15,20,25,30,35 | 0,10,20,25,35 |
| O2 | 0,5,10,20,25,35 | 0,20,25,35 |
| O3 | O1 schedule; delayed availability | O1 schedule; delayed availability |

For O1/O2, standardized observation error is $0.5\,b_{ij}+0.5\,v_{\mathrm{it}}+\sqrt{0.5}\,\varepsilon_{itj}$. Channel-specific noise scales and native/log coordinates are specified in Table S4. O3 substitutes a variance-standardized t5 independent component and multiplies the combined error by 1.5. Native channels add scaled error; log channels use the log1p/expm1 transform and nonnegative truncation. O0 supplies noiseless measurement-generating values annually through year 35.

The subject–channel component is a persistent offset, the visit component is shared across channels at that visit, and the remaining component is observation-specific. Before their specified reuse, the O1/O2 components are independent standard-normal draws. For a native-scale channel, recorded Y equals its noiseless mapped value plus the channel scale times this error. For a log-scale channel, recorded Y=max{exp[log(1+Y_noiseless) +s_log·error]−1,0}. Consequently, the noiseless mapping is not necessarily the arithmetic mean of recorded measurements. O3 retains the persistent and shared-visit components; its independent t5 component is rescaled to unit variance.

Postbaseline O1/O2 missingness probability is 0.05, with a 0.01 dropout hazard at scheduled visits. O3 uses missing probability $0.05+0.20\,sigmoid\left(5\left(\overline{X}-0.5\right)\right)$ and dropout hazard $0.02+0.08\,\overline{X}$. Dropout persists. Scheduled baseline values are retained by the baseline-completeness rule. O3 delays laboratory results by 7/365.25 years and imaging by 30/365.25 years; CAC remains in the laboratory reporting group. Other channels have no added delay. Eligibility requires both acquisition and availability by the cutoff.

Here X is mean burden across four domains. Dropout is a persistent visit-level hazard among individuals still observed. Both acquisition and availability must occur on or before the cutoff: delayed year-20 results are unavailable at that cutoff, and later results cannot complete earlier histories.

Whole-window gap limits are one year for O0; five years for core and ten for enriched channels in O1/O3; and ten years in O2. Unsupported windows stay unavailable. Endpoint-only trend has its explicitly allowed support rule and does not certify support for the whole-window mean. Current values have maximum ages of five years for P0/P1 channels and ten for P2 additions. Per-channel support comprises finite pre-cutoff count, latest age and maximum gap. Training medians and missing indicators belong to fitted preprocessing, not fabricated measurements.

#### S2.2 All 35 measurement mappings

P0, P1 and P2 contain six, nineteen and thirty-five channels, respectively, with nested membership. “First panel” identifies the smallest containing panel. Native scales use stated units; log scales use log(1+value). All mappings and scales are synthetic assumptions. Indices a/g/l/h denote domains and a_now denotes current age.

**Table S3. Complete principal measurement dictionary, first panel, units and reporting group.**

| ID | Measurement | First panel | Units | Reporting group |
|---|---|---|---|---|
| WC | Waist circumference | P0 | cm | anthro |
| SBP | Systolic blood pressure | P0 | mmHg | bp |
| DBP | Diastolic blood pressure | P0 | mmHg | bp |
| TG | Triglycerides | P0 | mg/dL | lab |
| HDL | HDL cholesterol | P0 | mg/dL | lab |
| FPG | Fasting glucose | P0 | mg/dL | lab |
| INS | Fasting insulin | P1 | uU/mL | lab |
| HBA1C | HbA1c | P1 | percent | lab |
| OGTT120 | Two-hour simulated glucose challenge response | P1 | mg/dL | lab |
| BMI | Body mass index | P1 | kg/m2 | anthro |
| FAT | Body fat percentage | P1 | percent | anthro |
| LDL | LDL cholesterol | P1 | mg/dL | lab |
| APOB | ApoB | P1 | mg/dL | lab |
| APOA1 | ApoA1 | P1 | mg/dL | lab |
| LPA | Lipoprotein(a) | P1 | mg/dL | lab |
| MET | Exercise capacity | P1 | MET | functional |
| LVMI_ECHO | Echo LV mass index | P1 | g/m2 | imaging |
| LVEF_ECHO | Echo LV ejection fraction | P1 | percent | imaging |
| CYS | Cystatin C | P1 | mg/L | lab |
| UACR | Urine albumin/creatinine ratio | P2 | mg/g | lab |
| LVMI_CMR | CMR LV mass index | P2 | g/m2 | imaging |
| LVEDVI | CMR LV end-diastolic volume index | P2 | mL/m2 | imaging |
| LVEF_CMR | CMR LV ejection fraction | P2 | percent | imaging |

| ID | Measurement | First panel | Units | Reporting group |
|---|---|---|---|---|
| RVEF | CMR RV ejection fraction | P2 | percent | imaging |
| RVEDVI | CMR RV end-diastolic volume index | P2 | mL/m2 | imaging |
| CAC | Coronary calcium | P2 | Agatston score | lab |
| IMT | Carotid intima-media thickness | P2 | mm | imaging |
| DIST | Carotid distensibility | P2 | 10^-3/kPa | functional |
| FMD | Flow-mediated dilation | P2 | percent | functional |
| ABI | Ankle-brachial index | P2 | ratio | functional |
| LIVER_HU | Liver CT attenuation | P2 | HU | imaging |
| NTBNP | NT-proBNP | P2 | pg/mL | lab |
| TROP | High-sensitivity troponin | P2 | ng/L | lab |
| LAVI | Left atrial volume index | P2 | mL/m2 | imaging |
| FFA | Free fatty acids | P2 | mmol/L | lab |

**Table S4. All 35 noiseless measurement mappings and channel-specific observation-error scales.**

| ID | Noiseless mapped value | Error scale | Error coordinate |
|---|---|---|---|
| WC | (74 if sex=F else 86) + 30*X_a | 1.0 | native_sd |
| SBP | 100+65*X_h | 4.0 | native_sd |
| DBP | 60+35*X_h | 3.0 | native_sd |
| TG | 70*exp(1.8*X_l) | 0.1 | log_sd |
| HDL | (65 if sex=F else 57)-25*X_l-5*X_a | 2.0 | native_sd |
| FPG | 80+100*X_g | 3.0 | native_sd |
| INS | 4*exp(1.6*X_a+1.2*X_g)*(.6+.4*Q_g) | 0.12 | log_sd |
| HBA1C | 4.6+3.4*X_g | 0.12 | native_sd |
| OGTT120 | 90+140*X_g+60*(1-Q_g) | 5.0 | native_sd |
| BMI | 19+17*X_a | 0.4 | native_sd |
| FAT | (18 if sex=F else 10)+30*X_a | 1.0 | native_sd |
| LDL | 70+110*X_l | 4.0 | native_sd |

| ID | Noiseless mapped value | Error scale | Error coordinate |
|---|---|---|---|
| APOB | 55+85*X_l | 3.0 | native_sd |
| APOA1 | 165-55*X_l | 4.0 | native_sd |
| LPA | LPA_i | 0.1 | log_sd |
| MET | 5+10*Q_h-3*X_h | 0.5 | native_sd |
| LVMI_ECHO | 55+45*X_h+55*M_h | 4.0 | native_sd |
| LVEF_ECHO | 70-12*X_h-20*(1-Q_h) | 2.5 | native_sd |
| CYS | .6+.6*X_h+.5*(1-Q_h) | 0.06 | log_sd |
| UACR | 3*exp(3*X_h+2*M_h) | 0.15 | log_sd |
| LVMI_CMR | 55+45*X_h+55*M_h | 2.0 | native_sd |
| LVEDVI | 55+25*X_h+20*M_h | 2.5 | native_sd |
| LVEF_CMR | 70-12*X_h-20*(1-Q_h) | 1.5 | native_sd |
| RVEF | 65-10*X_h-15*(1-Q_h) | 2.0 | native_sd |
| RVEDVI | 60+25*X_h+15*M_h | 2.5 | native_sd |
| CAC | 0 if a_now<35 else (a_now-35)*max(X_l-.15,0)*8+200*M_l | 0.2 | log_sd |
| IMT | .45+.35*X_h+.40*M_h | 0.03 | native_sd |
| DIST | 30/(1+2*O_h+2*M_h+X_h) | 0.1 | log_sd |
| FMD | 2+8*Q_h-3*X_h | 0.7 | native_sd |
| ABI | 1.15-.20*X_h-.20*M_h | 0.03 | native_sd |
| LIVER_HU | 65-35*X_a-10*X_l | 2.0 | native_sd |
| NTBNP | 20*exp(2.5*X_h+1.5*M_h) | 0.15 | log_sd |
| TROP | 2*exp(1.5*X_h+2*M_h) | 0.12 | log_sd |
| LAVI | 18+20*O_h+15*M_h+10*X_h | 2.0 | native_sd |
| FFA | .20+.60*X_a+.25*(1-Q_a) | 0.1 | log_sd |

##### S2.3 Restricted virtual research measurements

P3 is a restricted extension of P2 for G2/G3 under O1. Virtual input–response curves and recovery episodes are acquired at years 10 and 19. Recovery is evaluated on a local diagnostic copy and does not alter the long-term trajectory. The descriptors use measured probe responses, not true Q or O as predictor features. These probes were designed within the simulation and do not establish practical assay feasibility or clinical validity.

**Table S5. Virtual curve and recovery measurement laws, noise and support rules.**

| Frozen field | Definition |
|---|---|
| scope | G2/G3 O1 P3; not a causal clinical intervention protocol |
| Curve noise | u grid [0,.25,.50,.75,1]; response f(u)=sigmoid((u-(.35+.35*O_h))/(.08+.12*(1-Q_h))); Additive zero-mean Gaussian response noise with standard deviation 0.01, $\epsilon_{curve} \sim N(0, 0.01^2)$; reference O_h=.20,Q_h=.75; output normalized functional response. |
| recovery_episode | Local diagnostic copy with excursion D(s)=.20*exp(-(.4+Q_h)*s), s=[0,.25,.5,1,2,4] days, Additive zero-mean Gaussian response noise with standard deviation 0.005, $\epsilon_{recovery} \sim N(0, 0.005^2)$. No impact on main long-term trajectory, all history completed before cutoff. |
| features | ["MM2D-R03", "MM2D-R04", "MM2D-R05", "MM2D-O01", "MM2D-O02", "MM2D-O03"] |
| guards | Fixed input curve branches; target response .5; inverse interpolation only observed range; recovery resolution .015; t90 dwell 1 day; no use of true Q/O as learner feature. These measurements are deliberately designed virtual probes and cannot establish clinical assay validity. |

### S3. Representations, learning and complete retained Φ coordinates

#### S3.1 Conventional categorical and continuous inputs

The primary synthetic metabolic-syndrome comparator used waist circumference ≥102 cm for the conventional male category or ≥88 cm for the conventional female category; triglycerides ≥150 mg/dL or matching treatment; HDL cholesterol <40 mg/dL for the male category or <50 mg/dL for the female category, or specific low-HDL treatment; systolic blood pressure ≥130 mmHg or diastolic blood pressure ≥85 mmHg, or qualifying hypertension treatment; and fasting glucose ≥100 mg/dL or glucose-lowering treatment. Binary status required at least three of the five criteria, without a mandatory waist component. These were prespecified experimental comparator conventions, not new clinical recommendations. The separate O1/P0 sensitivity changed only the waist thresholds to ≥94 cm for the male category and ≥80 cm for the female category. All other component thresholds, treatment rules, handling of unknown criteria and the at-least-three-criteria rule remained unchanged; the sensitivity was not selected using held-out results.

Each criterion is true, false or unknown. With $c_{min}$ true criteria and $c_{max} = c_{min} + \# unknown$, binary status is positive if $c_{min} \geq 3$, negative if $c_{max} < 3$, and otherwise indeterminate. Count inputs retain the lower/upper pair. Blood pressure is negative only when both measured components are finite and low and no qualifying treatment is present. Classification never uses imputed values.

Current continuous input uses each channel's latest eligible value. Current-plus-baseline adds the recorded baseline values without adding intermediate observations. Raw history contains the declared pre-cutoff source-visit values and their missingness, without future or latent columns. Glycaemic history uses fasting-glucose history and adds insulin, HbA1c and simulated two-hour glucose only when the panel contains them. It does not borrow unmeasured channels or supply a true etiological label. A categorical label is a coarsening of its stated input; comparison of fitted predictors also depends on auxiliary access and fitting, so fitted loss differences do not isolate learner-independent information loss.

#### S3.2 Longitudinal operators and support

For an observed supported curve $y(s)$, orientation $a_j$ (also $o_j$) and reference $r_j$, mean departure is $W^{-1} \int_{t-W}^{t} a_j [y(s) - r_j] ds$. Accumulated excess integrates the positive part over ten years. Endpoint trend is $[y(t) - y(t-5)]/5$. Shape is $W^{-1} \int_{t-W}^{t} |y(s) - l(s)| / s_{j,baseline} ds$, where $l$ is the endpoint chord, $s_{j,baseline}$ is the training-baseline measurement SD, and an interior knot is required. Piecewise-linear operator interpolation is part of these feature definitions. Conventional references use the declared thresholds, HDL orientation is negative, and other references use the fitted baseline median.

The operator identifiers used in the retained-coordinate tables are B02 for time-mean oriented departure, B03 for accumulated positive excess, and B04 for endpoint trend. W5 and W10 mean five- and ten-year windows ending at the prediction cutoff. CURRENT means the latest eligible raw value. Means use time-weighted trapezoidal integration of eligible piecewise-linear observations. Positive-part integration inserts zero-crossing knots before integration; missing history is not zero exposure. Endpoint trend uses eligible bracketing observations without extrapolation. Shape uses exact piecewise-linear integration relative to the endpoint chord, including crossings, and requires at least one interior observed knot. A straight observed path has zero shape.

B02 has native measurement units, B03 native units×time, and B04 native units/time. Longitudinal windows use years; recovery episodes use days. Generic dictionary unit labels do not perform a time conversion. Conventional references use declared thresholds, with negative HDL orientation; other references and shape scales use fitting-population baseline medians and standard deviations. These are descriptive references, not universal healthy states.

All observations consumed by an operator must have been both acquired and available by the queried cutoff. Fixed windows must be complete under the specified maximum-gap rule; an unsupported window remains unavailable rather than being shortened. The endpoint-trend rule can be satisfied when whole-window support is not, and does not validate a mean or accumulated-excess calculation. Fitting medians, missingness indicators and support metadata are preprocessing objects rather than invented measurements.

##### S3.3 Research-probe operators

The six probe operators retain finite-support and censoring rules. Source-removal metadata must identify the actual input regime. Recovery to a prior baseline is not necessarily recovery to health; every consumed observation must be available by cutoff.

For the virtual recovery episodes, the specified baseline was zero, the recovery anchor was day zero, and the observation horizon was four days. Observations were made at $s=0,0.25,0.5,1,2,4$ days. The excursion amplitude used by R04 and R05 was the observed initial absolute excursion, $D_{peak}=|D_{obs}(0)|$. An initial excursion no greater than 0.015 left these two quantities unavailable. The input-response sweep used $u=0,0.25,0.5,0.75,1$, and observed responses were not clipped to the unit interval. Each complete curve-and-recovery episode became available at the episode year plus $4/365.25$ years.

For O01, the entire observed five-point response curve had to be monotone, and the target response 0.5 had to lie within the response ranges of both the observed and reference curves. Both inverses used piecewise-linear interpolation on the sampled curves, without extrapolation. The noiseless reference used $O_h=0.20$ and $Q_h=0.75$ on the same five-point input grid. Nonmonotone curves and out-of-range targets remained unavailable. A non-unique inverse retained its interval status but supplied no scalar predictor value. O03 was calculated only when both episode-specific shifts were finite scalar values.

For R05, the procedure identified the last sampled observation outside the recovery band and linearly interpolated the subsequent inward crossing. A crossing was retained only when at least one day remained between that crossing and the four-day endpoint. If the final observation remained outside the band, the result retained RIGHT_CENSORED status with a missing numerical predictor value. If the remaining dwell was shorter than one day, it retained INSUFFICIENT_DWELL status with a missing numerical value. The procedure used the complete fixed observation grid without an additional maximum-gap exclusion.

**Table S6a. Implemented signed residual (R03).**

| Frozen field | Definition |
|---|---|
| symbol | $r_4$ |
| formula | $$r_4=D_{obs}(4)$$ |
| algorithm | Use the observed day-four excursion relative to the specified virtual baseline of zero. Preserve its sign. R03 does not require the amplitude-resolution threshold used for R04 and R05. |

| Frozen field | Definition |
|---|---|
| units | normalised response |
| required_data | Specified baseline of zero and the observed day-four recovery value; episode timing and input-regime metadata. |
| availability_rule | All consumed observations acquired AND available by cutoff; all selected windows end by cutoff; external reference/configuration frozen independently of held-out outcomes. |
| limitations | Residual is not automatically memory, damage or feedback; ongoing internal burden can remain after external source removal. |

**Table S6b. Implemented recovered fraction (R04).**

| Frozen field | Definition |
|---|---|
| symbol | $\rho_4$ |
| formula | $\rho_4 = 1 - \frac{\left\|D_{obs}(4)\right\|}{D_{peak}}, \quad D_{peak} = \left\|D_{obs}(0)\right\| > 0.015$ |
| algorithm | Use the absolute initial observed excursion as the amplitude. An amplitude at or below 0.015 gives NO_RESOLVABLE_EXCURSION and a missing numerical value. Return the raw recovered fraction without clipping. |
| units | dimensionless |
| required_data | Specified baseline of zero, observed recovery values at days zero and four, and the fixed amplitude threshold of 0.015. |
| availability_rule | All consumed observations acquired AND available by cutoff; all selected windows end by cutoff; external reference/configuration frozen independently of held-out outcomes. |
| limitations | Undefined for an initial amplitude at or below the resolution threshold; may be negative when the late excursion exceeds the initial observed amplitude. Return to baseline need not mean return to health. |

**Table S6c. Implemented sustained recovery-band time (R05).**

| Frozen field | Definition |
|---|---|
| symbol | $t_{0.90}$ |
| formula | $t_{0.90} = inf\left\{t \in [0,4] : \left\|D_{obs}(s)\right\| \leq 0.1\, D_{peak} \quad \forall s \in [t,4]\right\}$ |

| Frozen field | Definition |
|---|---|
| algorithm | Use the specified baseline and anchor of zero and the four-day horizon. On the piecewise-linear observed trajectory, identify the last sampled point outside the band and interpolate the next inward crossing. Require at least one day of terminal dwell. A final point outside the band gives RIGHT_CENSORED; shorter dwell gives INSUFFICIENT_DWELL. Both statuses retain a missing numerical predictor value. An initial amplitude at or below 0.015 gives NO_RESOLVABLE_EXCURSION. |
| units | day |
| required_data | Observed recovery at days 0, 0.25, 0.5, 1, 2 and 4; initial absolute amplitude greater than 0.015; recovery fraction 0.90; band ±0.1 times that amplitude; minimum terminal dwell of one day. No additional maximum-gap exclusion was applied. |
| availability_rule | All consumed observations acquired AND available by cutoff; all selected windows end by cutoff; external reference/configuration frozen independently of held-out outcomes. |
| limitations | Finite-window observation only; right-censoring or insufficient dwell does not establish failed recovery. The result cannot establish sustained recovery beyond the four-day horizon. |

**Table S6d. Implemented input shift at matched output (O01).**

| Frozen field | Definition |
|---|---|
| symbol | $\Delta_u$ |
| formula | $\Delta_u = f_{obs}^{-1}(0.5) - f_{ref}^{-1}(0.5)$ |
| algorithm | Require the complete observed five-point curve to be monotone increasing or decreasing and the target 0.5 to lie within both sampled curve ranges. Invert by piecewise-linear interpolation without extrapolation or post hoc branch selection. A non-unique inverse can retain interval status but supplies no scalar predictor value. Nonmonotone curves and out-of-range targets remain unavailable. |
| units | normalised input |
| required_data | Observed and sampled noiseless-reference responses at inputs 0, 0.25, 0.5, 0.75 and 1; common target 0.5; fixed reference settings; episode acquisition and availability times. |
| availability_rule | All consumed observations acquired AND available by cutoff; all selected windows end by cutoff; external reference/configuration frozen independently of held-out outcomes. |
| limitations | No extrapolation or forced unique inverse. A curve shift is descriptive and does not by itself prove an internal setpoint mechanism. |

**Table S6e. Implemented response-curve secant (O02).**

| Frozen field | Definition |
|---|---|
| symbol | $S_u$ |

| Frozen field | Definition |
|---|---|
| formula | $$S_u = \frac{f_{obs}(0.75) - f_{obs}(0.25)}{0.50}$$ |
| algorithm | Use the observed responses at the declared input-grid points 0.25 and 0.75 and divide their difference by 0.50. This secant does not require the inverse-monotonicity guard used for O01. |
| units | normalised response per normalised input |
| required_data | Eligible observed curve values at inputs 0.25 and 0.75, with their episode timing and support. |
| availability_rule | All consumed observations acquired AND available by cutoff; all selected windows end by cutoff; external reference/configuration frozen independently of held-out outcomes. |
| limitations | Not a molecular gain or an infinitesimal derivative; interpretation is specific to the declared input interval and direction. |

**Table S6f. Implemented operating-shift change rate (O03).**

| Frozen field | Definition |
|---|---|
| symbol | $v_O$ |
| formula | $$v_O = \frac{\Delta_u(19) - \Delta_u(10)}{9}$$ |
| algorithm | Repeat O01 at target 0.5 using the same reference at years 10 and 19. Calculate the rate only when both shifts are finite scalar values. Otherwise retain an unavailable status and a missing numerical predictor value; interval-valued shifts do not yield a rate feature. |
| units | normalised input per year for the present study |
| required_data | Eligible comparable curve assessments at years 10 and 19, both supplying finite scalar shifts, with their acquisition and availability times. |
| availability_rule | All consumed observations acquired AND available by cutoff; all selected windows end by cutoff; external reference/configuration frozen independently of held-out outcomes. |
| limitations | Not evidence of restoration unless direction relative to health is independently established; same-subject reference uncertainty is correlated. |

#### S3.4 Development selection and the full 56-value Φ roster

Development ranked descriptors by absolute standardized Ridge coefficients averaged across specified outputs, tasks and cohorts, resolving ties by stable identifier. Missingness indicators travelled with descriptors but did not determine biological-value rank. The bounded menu included base/shape designs and 8, 16, 32, 64 or all values. Selection preferred the smallest candidate within 2% of raw-history average loss and within 10% in every scenario/channel guard. Otherwise it retained minimum loss, breaking ties toward fewer values, with an explicit no-adequate-compact-candidate status. No exhaustive subset optimality is claimed.

All eight P0, sixteen P1 and thirty-two P2 retained values are listed in order. Identifiers are coordinate keys. Baseline, context, missingness and support fields are additional, so displayed dimension is neither complete predictor size nor measurement count.

**Table S7a. All eight retained P0 Φ values.**

| Order | Exact feature ID |
|---|---|
| 1 | HDL__CURRENT |
| 2 | WC__CURRENT |
| 3 | FPG__CURRENT |
| 4 | TG__CURRENT |
| 5 | SBP__CURRENT |
| 6 | DBP__CURRENT |
| 7 | SBP__MM2D-B02_W10 |
| 8 | DBP__MM2D-B02_W10 |

**Table S7b. All sixteen retained P1 Φ values.**

| Order | Exact feature ID |
|---|---|
| 1 | WC__CURRENT |
| 2 | HDL__CURRENT |
| 3 | SBP__CURRENT |
| 4 | DBP__CURRENT |
| 5 | FAT__CURRENT |
| 6 | TG__CURRENT |
| 7 | FPG__CURRENT |
| 8 | DBP__MM2D-B02_W10 |
| 9 | BMI__CURRENT |
| 10 | SBP__MM2D-B02_W5 |
| 11 | DBP__MM2D-B02_W5 |
| 12 | FAT__MM2D-B02_W5 |
| 13 | SBP__MM2D-B02_W10 |
| 14 | OGTT120__CURRENT |

| Order | Exact feature ID |
|---|---|
| 15 | MET__CURRENT |
| 16 | TG__MM2D-B02_W5 |

**Table S7c. All thirty-two retained P2 Φ values.**

| Order | Exact feature ID |
|---|---|
| 1 | WC__CURRENT |
| 2 | HDL__CURRENT |
| 3 | FAT__CURRENT |
| 4 | FPG__CURRENT |
| 5 | FAT__MM2D-B02_W5 |
| 6 | TG__CURRENT |
| 7 | FAT__MM2D-B02_W10 |
| 8 | DBP__CURRENT |
| 9 | OGTT120__CURRENT |
| 10 | SBP__CURRENT |
| 11 | DBP__MM2D-B02_W10 |
| 12 | TG__MM2D-B02_W5 |
| 13 | FPG__MM2D-B02_W5 |
| 14 | BMI__CURRENT |
| 15 | TG__MM2D-B02_W10 |
| 16 | HDL__MM2D-B02_W5 |
| 17 | DBP__MM2D-B02_W5 |
| 18 | HBA1C__CURRENT |
| 19 | SBP__MM2D-B02_W5 |
| 20 | OGTT120__MM2D-B02_W5 |
| 21 | SBP__MM2D-B02_W10 |
| 22 | HDL__MM2D-B02_W10 |
| 23 | LIVER_HU__CURRENT |

| Order | Exact feature ID |
| --- | --- |
| 24 | FPG__MM2D-B02_W10 |
| 25 | APOB__CURRENT |
| 26 | BMI__MM2D-B02_W5 |
| 27 | OGTT120__MM2D-B02_W10 |
| 28 | FPG__MM2D-B03_W10 |
| 29 | HBA1C__MM2D-B02_W10 |
| 30 | LDL__CURRENT |
| 31 | APOA1__CURRENT |
| 32 | HBA1C__MM2D-B02_W5 |

##### S3.5 Selected and learned alternatives, preprocessing and tuning

The selected common representations retain 48, 215 and 311 values for P0, P1 and P2. The learned common representations use 8, 12 and 12 coordinates from respective upstream pools of 72, 215 and 311 values. Task-specific specialists were selected for their assigned continuation or low-load task, whereas common recipes served both absolute tasks. Selection did not inspect an individual held-out outcome. Table S8 records the locked recipe identities and their development qualification; complete ordered upstream lists for these alternatives are not supplied with this supplement.

**Table S8. Locked selected, learned and task-specific recipes. Historical T2_WITHDRAW identifiers denote the specified low-load reassignment.**

| Recipe | Task | Candidate | Dim. | Meaning | Value-source channels | Development lock status |
| --- | --- | --- | --- | --- | --- | --- |
| P0_X_A_COMMON | Both absolute tasks | A_FWD48 | 48 | Selected values | 6 | Point criterion passed; not compact |
| P0_X_B_COMMON | Both absolute tasks | B_RSV08 | 8 | Learned coordinates; nuisance excluded | 6 | No adequate compact candidate; best tested locked |
| P0_X_SPECIALIST_T1 | Continuation, absolute | A_FWD12 | 12 | Selected values | 6 | No adequate compact candidate; best tested locked |
| P0_X_SPECIALIST_T2 | Low-load, absolute | A_FWD48 | 48 | Selected values | 6 | Point criterion passed; not compact |
| P1_X_A_COMMON | Both absolute tasks | A_ALL | 215 | Selected values | 19 | No adequate compact candidate; best tested locked |

| Recipe | Task | Candidate | Dim. | Meaning | Value-source channels | Development lock status |
|---|---|---|---|---|---|---|
| P1_X_B_COMMON | Both absolute tasks | B_RSV12 | 12 | Learned coordinates; nuisance excluded | 19 | No adequate compact candidate; best tested locked |
| P1_X_SPECIALIST_T1 | Continuation, absolute | A_FWD12 | 12 | Selected values | 6 | No adequate compact candidate; best tested locked |
| P1_X_SPECIALIST_T2 | Low-load, absolute | A_ALL | 215 | Selected values | 19 | No adequate compact candidate; best tested locked |
| P2_X_A_COMMON | Both absolute tasks | A_ALL | 311 | Selected values | 35 | No adequate compact candidate; best tested locked |
| P2_X_B_COMMON | Both absolute tasks | B_RSV12 | 12 | Learned coordinates; nuisance excluded | 35 | No adequate compact candidate; best tested locked |
| P2_X_SPECIALIST_T1 | Continuation, absolute | A_FWD12 | 12 | Selected values | 7 | No adequate compact candidate; best tested locked |
| P2_X_SPECIALIST_T2 | Low-load, absolute | A_ALL | 311 | Selected values | 35 | Point criterion passed; not compact |

Locks fix identities, construction and dimension while permitting prescribed cohort-specific fitting of medians, scales, centres and projections. Preprocessing uses training data for validation and training-plus-validation for fixed-configuration refitting. Missingness remains distinguished from measurement. Unsupported columns are omitted without feature substitution; test outcomes cannot revise preprocessing, representation or historical maps.

Columns with no finite observations in the fitting population were omitted without replacement. Remaining missing entries were replaced by their column's fitting-population median. Supervised learner preprocessing appended one binary missingness indicator for every retained input column. The augmented matrix was centred using fitting-population means and scaled using population standard deviations, with divisor $n$; a zero standard deviation was replaced by one. The fitted column mask, medians, means and scales were reused on the corresponding validation or test records. Training data fitted the validation transforms, and training plus validation fitted the final transforms after configuration selection. Infinite feature values were invalid inputs rather than missing observations.

Cluster-map preprocessing used the same omission, imputation, centring and scaling steps without appended missingness indicators. The embedding's value projection also excluded appended indicators within its coordinate block; raw-value missingness and fixed auxiliary fields formed a separate nuisance block outside the locked coordinate count.

Each embedding teacher uses twelve absolute-outcome columns: six observed year-25 channels in each branch, excluding test individuals. Multioutput Ridge with penalty 1 fits each output on its finite permitted targets, standardized by their fitting-population mean/SD. Value and separate nuisance blocks have fitted preprocessing. Supported value-input coefficients supply a deterministic SVD basis; the representation projects preprocessed values onto its first locked k directions. Dimensions are 2, 4, 6, 8 or 12; inadequate rank makes a locked dimension unavailable. Missingness and fixed auxiliary fields remain outside k. This supervised linear embedding is neither neural nor unsupervised PCA, and coefficient rank is not biological dimension.

Ridge penalties are $\left(10^{-4}, 10^{-3}, 10^{-2}, 0.1, 1, 10, 100, 1000, 10000\right)$, with unpenalized intercept. Validation uses equal-output standardized MSE separately by task; ties within $10^{-10}$ choose the larger penalty. Arms need not select the same penalty. Refit uses training plus validation with decisions fixed. The scoring SD remains the original training observed-target scale and is distinct from head preprocessing scales.

Boosting uses squared-error loss, learning rate 0.05, 200 iterations, early stopping disabled, maximum leaves in{7,15,31}, L2 in{0,1}, minimum leaf size 30 and maximum bins 255. The six-point grid is a bounded O1/P0/P2 sensitivity.

K-means uses K=2–6, k-means++ initialization, 20 starts, 300 maximum iterations and tolerance $10^{-4}$. Training silhouette uses a deterministic sample capped at 1,000; ties within $10^{-4}$ favor smaller K, and candidates with a cluster below 2% of training subjects are rejected. Gaussian mixtures use diagonal covariance, five starts, 300 iterations, tolerance $10^{-3}$, covariance regularization $10^{-6}$, and minimum training BIC. The exact historical Gaussian-mixture initialization and library provenance remain incomplete. Hard membership is one-hot, soft membership is the posterior vector. Maps are fitted on training data for validation, then refitted on training plus validation with selection decisions fixed; final maps are reused for test historical assignment.

Historical membership used the frozen current map; the implemented $3K+2$ representation is specified in S3.6. Outcome-selected K uses the same K menu jointly with the nine Ridge penalties under validation loss and remains a separate sensitivity. No label-map refitting to improve apparent longitudinal stability is allowed.

##### S3.6 Implemented information access and measurement accounting

The membership-history representation (KM_HISTORY) comprised $K$ indicators for membership at the first scheduled query, $K$ at the final scheduled query, $K$ time-weighted dwell fractions, one transition count and one coverage fraction, giving $3K+2$ values before the addition of context and support fields. The map fitted to current measurements at year 20 was held fixed across historical queries. An all-missing current input vector received the unknown label −1 and all-zero membership indicators. Only adjacent query intervals with valid labels at both endpoints contributed: the earlier label received the interval duration, and a transition was counted when the two valid labels differed. Dwell fractions and coverage were divided by the full twenty-year horizon. Endpoint indicators used the first and last scheduled queries, rather than the first and last successful assignments. Baseline-only clusters used a separately fitted baseline map.

In the direct P0 absolute-task comparison, Φ has eight values plus six raw baseline values, thirteen context and eighteen support fields. Categorical history has 3K+2 values and shared context/support but lacks those raw baseline values. This compares longitudinal strategies, not a strict information hierarchy. Residual analyses retain their baseline control.

In this direct P0 comparison, the thirteen shared context fields were baseline age in years; sex; smoking; activity; observed E1 and E2; the four current input components $U_a, U_g, U_l, U_h$; and indicators of blood-pressure, triglyceride and glucose treatment. Sex was coded zero for the conventional female category and one for the conventional male category under the simulation's threshold convention. Observed E1/E2 retained their missing values, and current input components incorporated the stipulated treatment adjustment.

Each of the six P0 channels contributed three support fields: the number of finite observations acquired and available by the cutoff, the age in years of the latest such observation, and the largest interval in years between successive such observations. With no observations, the count was zero and age and gap were missing; with one observation, the gap was missing.

**Table S9. Value-source, learned-coordinate and nominal input-field accounting. The last column concerns value-source channels, including embedding upstream inputs; it excludes additional acquisition requirements imposed by baseline and panel support fields.**

| Recipe | Source values | Learned coordinates | Nominal input-field count | Channels in value block |
|---|---|---|---|---|
| P0::ORIGINAL_PHI_LOCKED | 8 | 0 | 53 | 6 |
| P0::X_A_COMMON | 48 | 0 | 133 | 6 |
| P0::X_B_COMMON | 72 | 8 | 117 | 6 |
| P0::X_SPECIALIST_T1 | 12 | 0 | 61 | 6 |
| P0::X_SPECIALIST_T2 | 48 | 0 | 133 | 6 |
| P1::ORIGINAL_PHI_LOCKED | 16 | 0 | 121 | 10 |
| P1::X_A_COMMON | 215 | 0 | 519 | 19 |
| P1::X_B_COMMON | 215 | 12 | 316 | 19 |
| P1::X_SPECIALIST_T1 | 12 | 0 | 113 | 6 |
| P1::X_SPECIALIST_T2 | 215 | 0 | 519 | 19 |
| P2::ORIGINAL_PHI_LOCKED | 32 | 0 | 217 | 14 |
| P2::X_A_COMMON | 311 | 0 | 775 | 35 |
| P2::X_B_COMMON | 311 | 12 | 476 | 35 |
| P2::X_SPECIALIST_T1 | 12 | 0 | 177 | 7 |
| P2::X_SPECIALIST_T2 | 311 | 0 | 775 | 35 |

Nominal input-field counts comprise representation values or learned coordinates, their designated value-missingness fields, and raw baseline, context and support fields under the stated accounting convention. They exclude additional indicators introduced by downstream preprocessing and do not report measured post-preprocessing matrix widths. Unsupported-column omission and constant-column handling depend on the fitted population.

Actual nonconstant usable columns can vary by cohort. The 10- and 14-channel footprints in the P1/P2 Φ rows describe the selected value block; baseline and panel support fields can still require the full declared panel. Observation counts and visit occasions also depend on window and availability rules. An embedding's small display dimension does not erase its upstream measurement requirements. No cost model, minimum biological dimension or complete-predictor acquisition saving is established by this accounting.

### S4. Outcomes, statistical procedures and multiplicity

#### S4.1 Targets, scoring and paired units

Primary targets are absolute recorded year-25 WC, SBP, DBP, TG, HDL and FPG under each input branch, with equal output weight. Residuals subtract observed baseline and require baseline control. Training uses channelwise observed targets with missingness; main held-out scoring uses complete synthetic noisy year-25 ascertainment. Observed-available-subset, noiseless-reference and latent-state evaluations retain separate target definitions.

Squared error was divided by the observed training-target population variance (ddof=0). Nonfinite outcome scales or standard deviations at or below $10^{-8}$ were invalid; the confirmation pipeline did not accept scores computed using a substituted floor. This scoring scale is not recomputed on test or refit populations and differs from head-preprocessing scales. Native RMSE/MAE retain units where reported. Compared methods share held-out identities, target masks and scales. Within a fixed scenario, inference used twenty independent confirmation cohorts as paired units. The balanced overall adequacy statistic used twenty prespecified replicate-index blocks, each containing one cohort from every scenario. Individuals, channels, future branches and observation replicas were not counted as additional independent inferential units.

#### S4.2 Primary and specialist inference

The primary paired analysis comprised 264 prespecified standardized-loss contrasts: 72 original contrasts and 192 extension contrasts. Each contrast used 20 paired confirmation cohorts and a two-sided paired Student $t$ statistic with 19 degrees of freedom. The sign convention was candidate minus comparator; negative values therefore indicated lower predictive loss for the candidate. Pointwise 95% $t_{19}$ intervals were retained as pointwise intervals and were not reinterpreted as simultaneous confidence intervals.

Primary multiplicity was handled in two frozen families. HOLM_JOINT_264 applied Holm step-down family-wise error control at $\alpha = 0.05$ across all 264 primary hypotheses. A separately specified historical family, HOLM_HISTORICAL_72, applied the same procedure to the original 72 hypotheses only. The historical family was not selected from the joint-family outcome. Joint-family correction uses the original unadjusted hypothesis p-values, not values already adjusted in another family.

The prespecified specialist family contained 108 task-specific specialist-versus-comparator contrasts under scheduled noisy observation. An exact predictive tie required matching prediction-value hashes for all six output channels in every one of the twenty cohorts, together with an exactly zero twenty-value loss-difference vector. Verified ties retained zero estimate, sample standard deviation, standard error and interval endpoints; their raw p-value was one, without calculating an ordinary t statistic. They remained in the full Holm family. Eighteen comparisons met this rule and retained Holm-adjusted p-values of one; the remaining ninety used paired two-sided t inference with nineteen degrees of freedom. A zero standard deviation without verified prediction identity retained no verdict. An unavailable p-value occupied its registered multiplicity slot with a working value of one for adjustment, while its reported p-value, adjusted p-value and scientific verdict remained undefined.

Splitting the study into two papers does not split, shrink or reset these original multiplicity families. Each family-specific adjusted p-value is interpreted within its original family and estimand. A pointwise interval remains pointwise when displayed beside an adjusted p-value. The companion papers reuse the same relevant cohorts and results; their presentation is not new confirmation.

#### S4.3 Simultaneous adequacy

Adequacy was defined by one-sided simultaneous upper confidence bounds on relative predictive loss versus RAW_HISTORY, not by failure to reject a null hypothesis. The simultaneous family contained 2,850 frozen predicates and used a one-sided $t_{19}$ upper bound at

$$q=1-\frac{0.05}{2850}=0.9999824561403509.$$

For a fixed candidate representation, panel and observation condition, let $L^{A}_{cgbj}$ and $L^{H}_{cgbj}$ denote the candidate and available-history standardised channel losses. Here $c=1,\ldots,20$ indexes confirmation replicates, $g=0,\ldots,5$ indexes scenarios, $b=1,2$ indexes the two absolute future tasks, and $j=1,\ldots,6$ indexes outcome channels. Each replicate-index block contains one independently simulated cohort from each scenario.

For a common representation, the balanced losses and overall relative excess for replicate $c$ were

$$A_c=\frac{1}{72}\sum_{g=0}^{5}\sum_{b=1}^{2}\sum_{j=1}^{6}L^{A}_{cgbj},H_c=\frac{1}{72}\sum_{g=0}^{5}\sum_{b=1}^{2}\sum_{j=1}^{6}L^{H}_{cgbj},r_{c,overall}=\frac{A_c-H_c}{H_c}.$$

Thus, scenarios, tasks and channels were equally weighted when averaging losses, before forming the overall relative excess. Each scenario–task–channel guard used

$$r_{c,gbj}=\frac{L^{A}_{cgbj}-L^{H}_{cgbj}}{L^{H}_{cgbj}}.$$

For a task-specific specialist, $A_c$ and $H_c$ instead averaged the thirty-six scenario–channel losses for its assigned task, with the same subsequent relative-loss calculation.

For every predicate $l$, the estimate, sample variance, standard error and simultaneous upper bound were

$$\bar{r}_{\ell}=\frac{1}{20}\sum_{c=1}^{20}r_{c,\ell},\qquad s_{\ell}^{2}=\frac{1}{19}\sum_{c=1}^{20}\left(r_{c,\ell}-\bar{r}_{\ell}\right)^{2}.$$

$$SE_l=\frac{s_l}{\sqrt{20}},U_l=\bar{r}_l+t_{19,\,1-0.05/2850}SE_l.$$

Adequacy required $U_{overall}\le 0.02$ and every applicable guard bound to be no greater than $0.10$. The simultaneous family comprised $36\times 73+6\times 37=2{,}850$ predicates. Common groups jointly covered both future tasks; specialist groups covered their assigned task.

Eligibility required the complete prescribed loss array, finite candidate and reference losses, and every required reference channel loss greater than $10^{-12}$. Failure of any of these requirements produced no verdict. Missing terms were not zero-filled, omitted or used to renormalise weights. After constructing the ratios, an undefined predicate also produced no group verdict before the margin conditions were evaluated. A zero sample standard deviation retained no verdict unless the comparison was a verified same-prediction alias with an exactly zero twenty-value difference vector. That exception retained zero estimate, standard error and upper bound without calculating an ordinary $t$ statistic.

For the 36 common recipe groups, each recipe required one balanced-overall predicate and 72 guard predicates. Adequacy required the overall upper bound to be no greater than 2% and every guard upper bound to be no greater than 10%. For the six specialist groups, each group required one overall predicate and 36 generator-by-channel guards under the same 2%/10% margins. Undefined evidence propagated to NO_VERDICT according to the frozen rule rather than being forced into adequacy or inadequacy.

For each predicate, the upper bound is its mean relative-loss estimate plus the stated t quantile times its cohort standard error. The accounting is 36×73+6×37=2,850 predicates across forty-two joint decisions. Reference loss at or below 10^−12 makes relative loss undefined; absolute loss can still be reported, but this case cannot certify adequacy. A joint failure does not establish that each task individually fails or that every margin is exceeded. Failure to reject a difference in another family is not an adequacy certificate. An exact predictive tie does not establish mechanistic equivalence.

##### S4.4 Additional comparisons and descriptive analyses

The current-input versus current-cluster completion used twelve scenario-by-task comparisons under O1/P0 with the same Ridge learner, six-channel equal-weight loss and twenty paired cohorts. Its difference is CURRENT loss minus KM_CURRENT loss. Each nondegenerate difference vector uses a two-sided t19 test and pointwise 95% t19 interval, with a separate Holm correction over twelve hypotheses. The question was planned, but this inferential completion was specified after related results were known; it is supplementary and cannot be described as retrospectively preregistered or blinded. It leaves the original primary, specialist and adequacy families unchanged.

Later alternatives retain their distinct families: learner sensitivity, 2,592 hypotheses; hard/soft mixture membership, 1,944; membership history, 3,528; outcome-selected cluster count, 324; and glycaemic-history input, 7,920. The direct Φ–membership-history comparisons have separate seventy-two-hypothesis families for Ridge and boosting; their 1,440 cohort–channel pairs per learner reuse the same 120 scenario–cohort combinations. A different joint saved-comparator analysis has 468 hypotheses including its specified interactions; no learner interaction is inferred from the two direct families alone. These reporting families do not denote independent datasets.

Undefined support, constant or incomplete data, non-rejection and zero difference remain distinct states. Eighteen outcome-selected-cluster channel hypotheses remain without a verdict because three tasks in one G0 cohort were unavailable. Order20 was the internal identifier for a planned aggregation of future outcome variation within shared categories. The underlying within-category summaries used sample variances of held-out year-25 measurements among individuals sharing a year-20 category, calculated separately by cohort, future branch and measurement channel in native squared units; each group required at least ten finite observations. The original aggregate was insufficiently specified and was not estimated. A separately specified amendment was completed after the original analysis campaign and retained its status as a separate later analysis. Natural tracking, repeat-measure localization and controlled response use the descriptive procedures below and are not converted into new confirmatory hypothesis families. Complete scorecards and paired-cohort evidence are listed as outstanding assets in S7.

### S5. Observation sensitivity, natural tracking and controlled movement

#### S5.1 Fixed-biology replicas and separate reference anchors

The fixed-biology procedure covers 120 O1/P0 scenario–cohort cells, the source-ordered first 256 held-out individuals in each, thirty remeasurements, twelve fitted Ridge methods, two absolute tasks and six channels. Across the thirty current-observation replicas, latent biology, medication, earlier observed history, the fitted pipeline and the persistent subject–channel bias were held fixed. The visit error was redrawn for each replica and shared across its channels, while residual observation errors were redrawn for each channel and replica. Both saved year-25 references were held fixed across replicas. Qualification cells are included within the 120 cells rather than added to them. Matching and differing category strata can contain the same individual under different replicas.

For one subject, method, task and channel, let $p_r$ be the prediction from remeasurement $r \in \{1, \ldots, 30\}$. The finite-replica mean and spread are

$$\overline{p} = \frac{1}{30} \sum_{r=1}^{30} p_r, V_R = \frac{1}{30} \sum_{r=1}^{30} \left( p_r - \overline{p} \right)^2.$$

Let $y_{25,noisy}$ and $y_{25,noise\text{-}free}$ denote the saved noisy and noise-free year-25 future references. The two future-reference errors are

$$\frac{1}{30} \sum_{r=1}^{30} \left( p_r - y_{25,noisy} \right)^2, \frac{1}{30} \sum_{r=1}^{30} \left( p_r - y_{25,noise\text{-}free} \right)^2.$$

The clean-current-input comparison instead records $D_{clean20} = \left( \overline{p} - p_{clean20} \right)^2$, where $p_{clean20}$ is the matched clean-current-input prediction from the same fitted pipeline. The cohort displacement averages the already computed subject-level squares. It is not a square applied to a pooled prediction difference. Scenario means, observed minima and maxima summarize cohorts with equal weight.

The noise-free future reference is one realised future mapped value, not an innovation-averaged expectation or clinical gold standard. Changing reference leaves predictions and spread fixed. The clean-current anchor instead removes current error, including persistent subject–channel bias, in the same fitted pipeline, retaining earlier noisy history, biology, medication and forced-observed current availability. P0 retains P0 access.

Replica spread, future-reference error and clean-current displacement are distinct. The last averages individual squared mean-to-clean-prediction differences before cohort reduction; it is not signed drift or biological bias. MSE decomposes into spread plus squared mean-to-reference discrepancy, but clean-current displacement cannot replace that discrepancy without equality of the anchors. A thirty-view mean is not a validated single-visit predictor.

#### S5.2 Coordinate variance, pipeline localization and scales

Native quantities use variance divisor 30. Complete 30-vectors are required; undefined coordinates remain null. Exact constant finite vectors have mathematical variance 0, avoiding floating-point mean-rounding artifacts. Independent verification used math.fsum means and the pairwise squared-difference identity with denominator 900, as well as independent cohort means, scale application and correlations.

Localization separates current/history values, preprocessed values, missingness and learned coordinates. Invariance requires complete defined replica vectors. Historical invariance under current-only remeasurement is a fixed-input control, not whole-history reliability. Constant finite vectors have zero mathematical variance; missing coordinates/scales remain undefined.

Per-cell scales support coordinate-specific profiles, not a universal cross-representation metric. Final coordinates use matched frozen head scales; native embedding inputs use encoder scales. No replica-derived scale, borrowed output scale or epsilon denominator is introduced. Learned axes are cell-specific. Variance correlations describe covariation, not mediation, causation or a biological interpretation of ordinal axes.

##### S5.3 Natural tracking and unavailable histories

Natural tracking evaluates eligible observed Φ coordinates at years 10 and 20 using acquisition and availability restrictions at each queried time. Prediction-head nuisance fields at year 20 are not silently substituted at year 10. Signed and absolute coordinate changes retain native units. Latent change D uses the defined subject-change quantity and the training-split year-20 standard deviations with population divisor (ddof=0). The within-cell association is Spearman(D, absolute coordinate change) on finite pairs. Constant or insufficient pairs remain undefined; no Fisher transformation, p-value, confidence interval or whole-vector norm is introduced.

$$D_i = \sqrt{\frac{1}{4}\sum_{d=1}^{4}\left[\frac{X_{id}(20) - X_{id}(10)}{s_{d,20,train}}\right]^2}.$$

The 1,440 eligible repeated observation/panel cells contain 1,473,120 cell–individual records but only 122,760 unique cohort–individual identities. Across-cohort means, medians and observed ranges remain descriptive. Forty P3 cells are held because of time support, and selected/learned whole-vector representations requiring history after year 10 are not replaced with different coordinates. Controlled perturbations answer a different question and do not repair these unavailable natural-tracking comparisons.

##### S5.4 Controlled perturbations, lanes and numerical resolution

Controlled response starts from saved year-15 state. One of U_a, U_g, U_l and U_h is changed to U_d+α(0.80−U_d), with α=0.1, 0.2 or 0.3 and the other inputs unchanged. No clipping or effect-selected replacement is added; zero input headroom retains its own status. Monthly continuation uses absolute months 180–239 to reach year 20. Lane A uses one shared clean control per cell/individual, zero future process innovations and clean endpoint mapped values, retaining the original observed prehistory. Lane B reuses the saved stochastic continuation only after the all-256-individual zero-dose agreement gate, with the keyed historical innovations and paired replicas 1–3. The two lanes are not pooled. Observed history through year 19 stays fixed while the year-20 endpoint is replaced; this is not a fully re-observed counterfactual history.

For paired endpoint values $z_1, z_0$, numerical resolution requires $|z_1 - z_0| > 2\,atol + rtol(|z_1| + |z_0|)$, using frozen layer-specific tolerances. Smaller differences remain stored rather than zeroed. Direction uses the predeclared direction table; absent signs remain absent. Ordering uses signed orientation where prescribed and absolute displacement otherwise, with undefined and all-unresolved states taking precedence over strict/nondecreasing/mixed classification. Numerical support is not a statistical test.

**Table S10. Numerical replay tolerances and retained qualification margins. The replay-equality formula differs from the response-resolution inequality in the text.**

| Frozen field | Definition |
|---|---|
| formula | abs(a-b) <= atol + rtol*max(abs(a),abs(b)) |
| safety_factor | 16 |

| Frozen field | Definition |
| --- | --- |
| safety_factor_justification | Prospective four-bit reserve over all observed zero-dose routes and original 5115 population. It covers the observed 1.6-fold increase in maximum absolute state error after population expansion with substantial remaining room. It is a conservative operational margin, not a probabilistic confidence bound or guarantee for untested generators/hardware. |
| states | {"atol": 2.842170943040401e-14, "rtol": 1.1368683772161603e-13, "ULP_ceiling": 1024, "observed_max_absolute": 1.3322676295501878e-15, "observed_max_symmetric_relative": 5.888187796884539e-15, "observed_max_ULP": 47, "absolute_margin_factor": 21.333333333333332, "relative_margin_factor": 19.30761070184754, "ULP_margin_factor": 21.78723404255319} |
| representations | {"Phi8": {"atol": 3.637978807091713e-12, "rtol": 2.842170943040401e-14, "ULP_ceiling": null, "observed_max_absolute": 1.4210854715202004e-13, "observed_max_unit_floor_relative": 1.245153706476022e-15, "observed_max_ULP": 8, "absolute_margin_factor": 25.6, "relative_margin_factor": 22.82586421466138}, "XA48": {"atol": 7.275957614183426e-12, "rtol": 4.547473508864641e-13, "ULP_ceiling": null, "observed_max_absolute": 3.481659405224491e-13, "observed_max_unit_floor_relative": 2.842170943040401e-14, "observed_max_ULP": 3277, "absolute_margin_factor": 20.897959183673468, "relative_margin_factor": 16.0}, "XB72_native": {"atol": 7.275957614183426e-12, "rtol": 4.547473508864641e-13, "ULP_ceiling": null, "observed_max_absolute": 3.481659405224491e-13, "observed_max_unit_floor_relative": 2.842170943040401e-14, "observed_max_ULP": 3277, "absolute_margin_factor": 20.897959183673468, "relative_margin_factor": 16.0}, "XB8_learned": {"atol": 1.1368683772161603e-13, "rtol": 5.684341886080802e-14, "ULP_ceiling": null, "observed_max_absolute": 4.440892098500626e-15, "observed_max_unit_floor_relative": 2.886579864025407e-15, "observed_max_ULP": 832, "absolute_margin_factor": 25.6, "relative_margin_factor": 19.692307692307693}} |

Lane-B signal relative to measurement variation is R=|perturbed−control|/sqrt(V), using the exact individual–coordinate variance V from the matched thirty-remeasurement reference. R is defined only for positive finite V and finite response. Zero measurement floors and undefined references retain separate statuses; neither an epsilon denominator nor infinity is substituted. Lane A has no matched R in this design. Smaller numerically unresolved differences remain stored unchanged rather than rounded to zero. Numerical resolution is an operational precision criterion, not a statistical significance test.

Direction is assessed only against the predeclared direction table; no learned-axis direction is invented. Weak expected signs allow zero headroom, saturation and fixed latest-observation slots. Ordering uses signed oriented response where a positive or negative expectation exists and absolute displacement otherwise. Undefined or all-unresolved states take precedence over strict, nondecreasing or mixed classifications. Frozen category maps give separate unknown, unchanged and changed statuses.

The category-silent fraction among numerically resolved coordinate changes, the category-silent fraction among changes exceeding one measurement SD, and the representation–category pair switch fraction have different denominators. Repeated response records are not individual responder rates. Raw replica-specific quantities are reduced first to equal-individual cell summaries and then to equal-cohort scenario summaries. Means, medians, observed minima/maxima and defined/expected cohort counts are descriptive; no p-values or uncertainty intervals are added. The procedure evaluates frozen-representation responsiveness to a controlled synthetic perturbation, without a prediction head and without establishing therapeutic efficacy or human validity.

### S6. Post hoc monthly-kernel lumpability analysis

#### S6.1 Target and scope

This separate exploratory analysis concerns the G0 one-month scientific transition law with noiseless P0 mappings at year 1. It is post hoc and is not retrospectively preregistered. Strong lumpability is tested directly through destination-category transition probabilities at fixed context, rather than importing a finite-state theorem as a result for the continuous model. The monthly query does not alter O0's annual observation schedule.

Write the sufficient state as $x=(a,g,l,h)$, the adiposity, glycaemic, lipoprotein and haemodynamic burden coordinates. At a fixed starting context $c$ and $\delta=1/12$ year, define the transition probability $K_c(x,D)$ into destination set $D$ by the exact monthly generator. Let $c^+$ advance age/time by $\delta$ under the fixed input policy. The category kernel is

$$p_b(x;c)=K_c\left(x,\{x':g\left(h\left(x',c^+\right)\right)=b\}\right).$$

Here $h\left(x',c^+\right)$ is the noiseless observation map and $g(\cdot)$ the category map, distinguished by their arguments from the burden coordinates $h$ and $g$; $b$ denotes a destination category. Strong lumpability requires equality of every $p_b$ for every two admissible states in the same current partition block at that same context. An unequal pair directly contradicts this universal condition; no homogeneity or stationarity assumption is used.

G0 fixes Q=0.75, O=0.20 and M=C=0. External context includes sex category, starting age, input, treatment and every other fixed quantity required by the mapping or transition law. Destination age advances by one month. This context is held equal between paired states; the six noiseless P0 measurement mappings themselves contain no age term.

#### S6.2 Exact monthly law and positive-density support

For domain index $d\in\{a,g,l,h\}$, let $r=(.18,.22,.25,.30)$ in that coordinate order and $\rho_d=\exp(-r_d\delta)$. Write the clipped age term as

$$A_{age}=clip\left((age-18)/50,0,1\right).$$

Then the target is

$$T_d=clip\left(.05+.65U_d+.08\frac{\sum_{k\neq d}x_k}{3}+.04A_{age},.01,.95\right).$$

The source uses all old states simultaneously. The deterministic update $d_d$ and its logit $\mu_d$ are

$$d_d=T_d+\left(x_d-T_d\right)\rho_d,\mu_d=logit\left(d_d\right),$$

and the next burden satisfies

$$logit\left(X_d^{'}\right)=\mu_d+\beta C+\gamma E_d,$$

where

$$\beta=.06\sqrt{.3\delta},\gamma=.06\sqrt{.7\delta},$$

and $C, E_a, E_g, E_l, E_h$ are independent standard normal variables. Here $C$ denotes the shared Gaussian innovation, distinct from the fixed cumulative carrier; $E_d$ denotes the domain-specific innovation. Innovations are independent across months. Thus the logit covariance is

$$.0003\left(.7I+.3 1 1^T\right):$$

its diagonal is $.0003$, its off-diagonal entries are $.00009$, and its smallest eigenvalue is $.00021$. Here $I$ is the four-dimensional identity matrix and $1$ is the four-component all-ones vector. The future state has positive density throughout $\left(0,1\right)^4$. There is no final state clipping in the source law. This statement concerns the specified Gaussian law, not finite pseudorandom-number-generator (PRNG) bit-orbit support.

The untreated input domain $\left[0.02,.80\right]^4$ puts $T$ between $.063$ and $.69$, inside the clipping endpoints. The implementation nevertheless retains the original clip operation. The probability law integrates innovations rather than replaying a saved seed.

The five criteria reduce to the latent events below. Here a, g, l and h denote burden coordinates; the category and observation maps are distinguished by their arguments. Positive-density support is a property of the specified Gaussian mathematical law, not a claim about all bit patterns reachable by a finite pseudorandom generator.

**Table S11. Exact latent threshold events under false treatment flags.**

| Criterion | Source measurements | Exact latent event, false treatment flags |
|---|---|---|
| Waist | 74+30a F; 86+30a M | a>=w, w=7/15 F or8/15 M |
| TG | 70 exp(1.8l) | l>=tau=log(15/7)/1.8 |
| HDL | 65-25l-5a F; 57-25l-5a M | l>s(a), s(a)=.6-.2a F or.68-.2a M |
| BP | SBP=100+65h, DBP=60+35h | h>=min(6/13,5/7)=6/13 |
| Glucose | 80+100g | g>=.2 |

HDL equality is negative; the others use non-strict positive thresholds. Exact future equalities have Gaussian probability zero, while classifier quality-assurance (QA) checks still test these conventions. TG/HDL dependence and the BP OR rule are retained.

#### S6.3 Destination probabilities with retained dependence

In the integration below, $c$ is the realised shared Gaussian innovation, distinct from the fixed external context denoted $c$ above. Write $\Phi_N$ and $\phi_N$ for the standard-normal cumulative distribution function (CDF) and density; these are distinct from the longitudinal representation $\Phi$. Condition on $C=c$ and $E_a=e$, so

$$a=sigmoid\left(\mu_a+\beta c+\gamma e\right).$$

For the future lipoprotein burden $L'=X_l^{'}$, set

$$F_l\left(v;c\right)=\Phi_N\left(\frac{logit\left(v\right)-\mu_l-\beta c}{\gamma}\right).$$

The joint lipid criterion count has polynomial

$$q_0 + q_1 z + q_2 z^2,$$

where $z$ is the polynomial variable and

$$\begin{aligned} q_0 &= F_l\left(min\left(\tau, s\left(a\right)\right); c\right), \\ q_1 &= Pr\left[min\left(\tau, s\left(a\right)\right) < L' < max\left(\tau, s\left(a\right)\right) \mid c\right], \\ q_2 &= 1 - F_l\left(max\left(\tau, s\left(a\right)\right); c\right). \end{aligned}$$

Multiply by $z^{1\{a \geq w\}}$, where $1\{\cdot\}$ is the indicator, and by the two conditional Bernoulli polynomials for glucose and BP. Their conditional independence holds because their distinct $E_d$ are independent once $C$ is fixed. Integrating coefficients against $\phi_N(c)\phi_N(e)$ yields all six METSCOUNT probabilities. METSBIN sums counts 0–2 and 3–5. Five unconditional criterion marginals are never multiplied.

Nested adaptive Gauss–Kronrod integration splits the adiposity-specific integral at its waist crossing and at $s(a) = \tau$. The variables $C$ and $E_a$ are integrated on $[-12, 12]$; the other normals are integrated by full-tail CDFs. The omitted mass is

$$1 - \left[1 - 2\Phi_N(-12)\right]^2 \leq 4\Phi_N(-12),$$

with the bound approximately $7.11 \times 10^{-33}$. Stable log-CDF/survival interval subtraction protects cancellation. The frozen two-level convergence ladder is an error estimate, not validated interval arithmetic. Entries below that estimate are explicitly unresolved in absolute accuracy; any rounded/underflow zero is not an asserted structural zero.

#### S6.4 Admissible same-category probes

The numerical examples use the same conventional female-threshold context and no qualifying treatment flags. Query age is 23.95274049 years and continued input, in a/g/l/h order, is (0.60267921, 0.70079018, 0.43457133, 0.46753891). These displayed age/input values are rounded to eight decimal places; the destination probabilities were calculated from the original full-precision record. Exact numerical replay requires that record, which is not supplied here. This precision limitation does not affect the analytic inequality under the stated same-context assumptions.

The saved admissible context was selected before probability calculation. At baseline, input and burden share Gaussian draws, so arbitrary joint combinations cannot be justified by coordinate bounds alone. The constructed probes are placed at month 12. Conditional on the admissible fixed context and any earlier admissible state, the nonsingular independent last-month innovation supplies positive density on every interior open burden box; mixing over previous states preserves that support. The four vectors are constructed model queries, not observations from four individuals.

**Table S12. All four constructed G0 burden states; each has current criterion count two and negative binary status in the specified female context.**

| Probe | Adiposity | Glycaemic | Lipoprotein | Haemodynamic | Current count | Current binary |
|---|---|---|---|---|---|---|
| P01_A | 0.5 | 0.19 | 0.4 | 0.5 | 2 | 0 |
| P01_B | 0.51 | 0.195 | 0.41 | 0.51 | 2 | 0 |
| P02_A | 0.4 | 0.19 | 0.53 | 0.44 | 2 | 0 |

| Probe | Adiposity | Glycaemic | Lipoprotein | Haemodynamic | Current count | Current binary |
|---|---|---|---|---|---|---|
| P02_B | 0.41 | 0.195 | 0.54 | 0.45 | 2 | 0 |

Within each pair, B exceeds A in every burden coordinate. Pair 1 satisfies waist and blood-pressure criteria; pair 2 satisfies triglyceride and low-HDL criteria. Pair-1 category-preserving L∞ neighbourhood radii are 0.0025 for A and 0.00125 for B. These positive radii show that the example does not require current diagnostic-boundary states. They account for state endpoints and the waist, glucose, TG, BP and HDL boundaries; the HDL plane has L∞ Lipschitz constant 30 in its measurement units.

#### S6.5 Full analytic proof of unequal category kernels

For each registered pair $x_B > x_A$ in every coordinate, the deterministic drift satisfies $d_B > d_A$: its own-coordinate coefficient $\rho_d$ is positive and the other-coordinate coefficients are nonnegative. Therefore

$$\Delta_d = \mu_{Bd} - \mu_{Ad} > 0 .$$

Couple the two next states using the same innovations. Then

$$X'_{Bd} = sigmoid\left(logit\left(X'_{Ad}\right) + \Delta_d\right) > X'_{Ad} .$$

Each criterion, the count and the METSBIN-positive event are nondecreasing in burdens, so the next criterion counts satisfy $N_B \geq N_A$ pathwise.

Let $w$ be the waist latent threshold. Consider the following open rectangle in A's future state:

$a \in \left(sigmoid\left(logit\left(w\right) - \Delta_a\right), w\right)$;

$l \in \left(\tau, s\left(w\right)\right)$;

$g \in \left(.25, .30\right)$;

$h \in \left(.20, .30\right)$.

A has exactly two criteria there: TG and glucose. Waist and BP are negative, and since $a < w$ and $s\left(a\right) > s\left(w\right) > l$, HDL is negative. B necessarily has waist, TG and glucose positive and hence count at least 3. All intervals are nonempty. In particular, $\tau < 1/2$ because

$$\exp\left(.9\right) > 1 + .9 + \frac{.9^2}{2} > \frac{15}{7} ,$$

whereas $s_F\left(w\right) = 38/75 > 1/2$ and $s_M\left(w\right) = 43/75 > 1/2$. The Gaussian density is strictly positive on this rectangle's logit preimage, so this event has positive probability.

Consequently,

$$p_{positive}\left(B\right) > p_{positive}\left(A\right)$$

The strict inequality is analytic and independent of quadrature precision, underflow or the estimated numerical gap. The two six-component count-probability vectors for each pair must also differ, because their sums over counts 3–5 differ. Numerical effect sizes remain estimates even though the inequality is proved.

This refutes strong lumpability for the examined one-month context-dependent kernel and both specified partitions. The noiseless full P0 map is invertible in the G0 current burdens; the coarsening loses within-category transition information. This does not infer annual-process non-lumpability, non-Markovianity for the particular original cohort distribution, five-year forecast superiority, clinical biological memory, stationarity, or a human metabolic-syndrome mechanism. No practical clinical importance threshold or population-average effect is introduced.

##### S6.6 Complete numerical destination estimates

The numerical tables retain all four probes, all six count destinations and both binary destinations. “Below resolution” means the reported probability is smaller than the numerical convergence estimate; it is not a structural zero. The error estimate is not a certified enclosure. The analytic proof establishes strict inequality independently of these estimates, while the magnitude of each numerical difference remains an estimated model quantity.

**Table S13. Complete destination probabilities and numerical convergence estimates for every constructed probe.**

| Probe | Partition | Destination | Probability | Error estimate | Below resolution |
|---|---|---|---|---|---|
| P01_A | METSCOUNT | 0 | 3.1234850104499935e-24 | 2.9929655229111635e-11 | True |
| P01_A | METSCOUNT | 1 | 1.2231868804029113e-14 | 2.9929655229111635e-11 | True |
| P01_A | METSCOUNT | 2 | 0.8984291454399391 | 2.9929655229111635e-11 | False |
| P01_A | METSCOUNT | 3 | 0.10157085116794214 | 2.9929655229111635e-11 | False |
| P01_A | METSCOUNT | 4 | 3.3921067212938346e-09 | 2.9929655229111635e-11 | False |
| P01_A | METSCOUNT | 5 | 5.443098238805679e-108 | 2.9929655229111635e-11 | True |
| P01_A | METSBIN | 0 | 0.8984291454399513 | 8.97889656873349e-11 | False |
| P01_A | METSBIN | 1 | 0.10157085456004886 | 8.97889656873349e-11 | False |
| P01_B | METSCOUNT | 0 | 3.472460252978325e-38 | 6.887497135836856e-12 | True |
| P01_B | METSCOUNT | 1 | 2.013950713825643e-23 | 6.887497135836856e-12 | True |
| P01_B | METSCOUNT | 2 | 0.30462729931147803 | 6.887497135836856e-12 | False |
| P01_B | METSCOUNT | 3 | 0.6950328455157115 | 6.887497135836856e-12 | False |
| P01_B | METSCOUNT | 4 | 0.0003398551728107082 | 6.887497135836856e-12 | False |
| P01_B | METSCOUNT | 5 | 1.8124879686843317e-83 | 6.887497135836856e-12 | True |
| P01_B | METSBIN | 0 | 0.30462729931147803 | 2.0662491407510567e-11 | False |
| P01_B | METSBIN | 1 | 0.6953727006885222 | 2.0662491407510567e-11 | False |
| P02_A | METSCOUNT | 0 | 2.1440181873872989e-128 | 6.347748005337131e-11 | True |
| P02_A | METSCOUNT | 1 | 0.06907032477930186 | 6.347748005337131e-11 | False |
| P02_A | METSCOUNT | 2 | 0.8318507575571824 | 6.347748005337131e-11 | False |
| P02_A | METSCOUNT | 3 | 0.09907888414797002 | 6.347748005337131e-11 | False |

| Probe | Partition | Destination | Probability | Error estimate | Below resolution |
|---|---|---|---|---|---|
| P02_A | METSCOUNT | 4 | 3.351554585955998e-08 | 6.347748005337131e-11 | False |
| P02_A | METSCOUNT | 5 | 1.5579999359030041e-55 | 6.347748005337131e-11 | True |
| P02_A | METSBIN | 0 | 0.9009210823364843 | 1.9043244016011393e-10 | False |
| P02_A | METSBIN | 1 | 0.09907891766351588 | 1.9043244016011393e-10 | False |
| P02_B | METSCOUNT | 0 | 7.69860582767465e-155 | 6.867868183381916e-12 | True |
| P02_B | METSCOUNT | 1 | 2.4956563912772407e-05 | 6.867868183381916e-12 | False |
| P02_B | METSCOUNT | 2 | 0.30638285415680677 | 6.867868183381916e-12 | False |
| P02_B | METSCOUNT | 3 | 0.6924640571716416 | 6.867868183381916e-12 | False |
| P02_B | METSCOUNT | 4 | 0.0011281321076390505 | 6.867868183381916e-12 | False |
| P02_B | METSCOUNT | 5 | 2.2028925195002962e-39 | 6.867868183381916e-12 | True |
| P02_B | METSBIN | 0 | 0.30640781072071954 | 2.0603604550145746e-11 | False |
| P02_B | METSBIN | 1 | 0.6935921892792807 | 2.0603604550145746e-11 | False |

**Table S14. Paired category-distribution differences and combined numerical error estimates.**

| ID | Partition | Origin | Maximum difference | Total variation | Combined error estimate |
|---|---|---|---|---|---|
| P01 | METSBIN | 0 | 0.5938018461284733 | 0.5938018461284733 | 1.1045145709484548e-10 |
| P01 | METSCOUNT | 2 | 0.5938018461284611 | 0.5938018461284733 | 3.681715236494849e-11 |
| P02 | METSBIN | 0 | 0.5945132716157648 | 0.5945132716157648 | 2.1103604471025969e-10 |
| P02 | METSCOUNT | 2 | 0.5933851730236716 | 0.5945132716157647 | 7.034534823675323e-11 |

#### S7. Software provenance, availability and reproducibility limits

The protocol prescribes NumPy Generator/PCG64 and fixed named streams. This prospective specification does not establish every exact library default used in execution. Historical Gaussian-mixture initialisation and library provenance remain incomplete, as recorded in the study documentation. Replay and qualification claims remain confined to the particular calculations and environments that were checked.

The controlled-response calculation records Python 3.13.13 (main, May 19 2026, 23:47:56) [GCC 14.2.0] and NumPy 2.3.5, with BLAS/OMP/MKL/NumExpr thread settings all 1. This is provenance for that calculation only; it does not establish a common environment for all analyses. Document-production software is separate from the scientific execution environments.

This supplement supplies the bounded generating assumptions, all 35 measurement mappings, all 56 retained Φ values, operator definitions, stated fitting procedures and the monthly analytic proof. It does not supply an executable reproduction package. The complete 264-comparison primary and 108-comparison specialist scorecards, full machine-readable forty-two-group adequacy scorecard with all fields, 2,850 predicate records and unrounded paired-cohort losses are unavailable with this supplement. Summarized decisions for all forty-two adequacy groups are presented in the second paper; this does not replace the full supporting scorecard and primitive evidence.

Also unavailable with this supplement are the complete ordered selected-feature and embedding-upstream rosters; fitted preprocessing, map and projection payloads; original random streams and exact execution configurations; complete repeat-measure, natural-tracking and perturbation banks with their joins; the full predeclared direction table; exact stored probe context; and study implementation code. The prose and selected numerical tables cannot substitute for these missing assets. Historical mixture initialization remains incompletely documented.

These omissions limit exact computational reproduction of the affected analyses. Additional study-specific materials may be requested from the corresponding authors, subject to ownership and release permissions; access is not guaranteed. The broader unpublished METLAS framework, confidential calibration resources and production implementation are not distributed with these papers.
